# Multi-Omics Reveals Activated Fibroblasts with Dual Functional Roles in Repair and Negative Barrier in the CD8^+^ T Cell Cytotoxic Niche for Lung Cancer Neoadjuvant Therapy

**DOI:** 10.64898/2026.01.14.699401

**Authors:** Ruihan Zhou, Chaoxin Xiao, Ping Zhou, Ouying Yan, Banglei Yin, Xiaohong Yao, Jiaxin Liu, Xuexue Wu, Wanting Hou, Yulin Wang, Huanhuan Wang, Rui Zhu, Zirui Wang, Leyi Yao, Xiaoying Li, Tongtong Xu, Fujun Cao, Na Xiao, Ke Cheng, Lili Jiang, Dan Cao, Chengjian Zhao

## Abstract

A major challenge in elucidating immune activation and tolerance is that single-omics technologies are inherently limited. Since single-cell transcriptomics lacks spatial information and spatial transcriptomics lacks resolution or depth, neither can adequately model the multidimensional tumor immune microenvironment (TiME) features. We present NICHE (Niche Integrated Cellular Heterogeneity Elucidator), an integrative multi-omics framework. By aligning physical cell-cell interactions with single-cell ligand-receptor (L-R) expression, NICHE leverages single-cell transcriptomics, spatial proteomics, and AI modeling to systematically decode the composition, spatial interactions, and communications within immune functional niches. To validate NICHE’s capabilities, we applied it to human tonsil tissue. The framework successfully identified key structural and functional units and deeply elucidated the cell-cell interactions and molecular crosstalk within them. Furthermore, these results were validated through orthogonal methods, including single-cell spatial transcriptomics and multiplexed immunofluorescence, which confirmed the accuracy and reproducibility of our analyses. In non-small cell lung cancer (NSCLC) patients undergoing neoadjuvant immunotherapy, CD8^+^ T cell cytotoxic niches mediate tumor clearance with variable outcomes, to define the cellular and molecular drivers behind this functional spectrum, we utilized NICHE to analyze the TiME from patients with complete versus partial pathological response, both pre- and post-treatment. We found that neoadjuvant immunotherapy may induce the activation of CXCL12^+^, NECTIN2^+^, POSTN^+^, and COL6A1^+^ fibroblasts through the activation of cytotoxic T cells. Paradoxically, these fibroblasts support tissue repair but also establish an immunosuppressive niche that protects residual tumor cells, ultimately attenuating treatment efficacy. Our work establishes a multidimensional framework for dissecting dynamic immune activity, with direct implications for understanding tumor immunology and identifying novel therapeutic targets.

## INTRODUCTION

The immune system serves as the body’s core barrier against pathogenic invasion, maintains homeostasis of the internal environment, and eliminates abnormal cells; it is thus essential for overall health. With advancing research, the immunological perspective has expanded beyond merely focusing on the defensive roles of immune cells to encompass their complex interactions with non-hematopoietic cells in tissue microenvironments.^1–4^ In this context, the concept of a “functional niche (FN)” has gradually taken shape. It describes a functional unit within a defined spatial range, jointly constructed by immune cells, non-immune cells, and local environmental factors (such as extracellular matrix, cytokines, vascular structure, etc.).^5–7^ This structural unit reflects the intrinsic, programmatic regulatory logic within tissues and determines how local immune homeostasis is maintained and how immune responses are orchestrated. Related studies often describe these structures as *“*neighborhoods,*” “*communities,*”* or *“*milieus,*”* emphasizing their importance in spatially dependent cell communication and functional regulation.^8–10^ The formation of functional domains relies on multilayered cellular interactions, encompassing not only direct cell-cell contacts but also paracrine signaling mediated by chemokines and cytokines. These interactions establish a coordinated immune response environment through intricate ligand-receptor (L-R) networks and signaling pathways. Representative L-R pairs, such as CD80-CD28, CD40-CD40L, TCR-MHC I/II, and PD-L1-PD-1 together^11–13^ with various cytokine pathways, including IL-12, IFN-γ, IL-10, and TGF-β,^14–16^ collectively drive the establishment of multicellular activation niches, enabling different immune cell types to exert complementary functions and ultimately achieve an efficient and well-orchestrated immune response.

In recent years, spatial proteomics technologies have advanced rapidly, including PhenoCycler-Fusion (formerly CODEX), imaging mass cytometry (IMC), and cyclic multiplexed tyramide signal amplification (CmTSA).^17–19^ Combined with deep learning-based algorithms such as StarDist^20^ and Cellpose^21^ for accurate cell segmentation, these approaches have made it possible to construct spatial proteomic maps with single-cell resolution across entire tissue sections, while preserving the in situ spatial context of each cell within the tissue architecture. Building upon this foundation, the integration of unsupervised clustering methods (e.g., K-means)^22,23^ and advanced modeling strategies such as graph neural networks (GNNs)^24^ enables researchers to identify and interpret multicellular functional structures that span the tissue landscape, thereby capturing intercellular interaction patterns and the hierarchical organization of the tissue microenvironment. This integrative strategy not only enhances the analytical depth of spatial proteomics data but also provides a systematic and fine-grained perspective for elucidating tissue development, pathological progression, and dynamic remodeling of the immune microenvironment.

However, the mere prediction of functional niches is insufficient to fully elucidate their biological essence. The integration of spatial and multi-omics technologies has opened new avenues for decoding the complexity of tumors and their microenvironments. By analyzing gene expression patterns and their spatial distributions within tumor tissue sections, researchers can not only map the localization and functional states of immune cells at single-cell resolution but also uncover their dynamic interactions with tumor and stromal cells. In particular, the combination of spatial proteomics, single-cell transcriptomics, and spatial transcriptomics has emerged as a central strategy for decoding tissue functional niches and their regulatory networks, providing a solid foundation for a systematic understanding of the multilayered mechanisms governing the tumor immune microenvironment. For example, in hepatocellular carcinoma (HCC), integrated single-cell RNA sequencing (scRNA-seq) and spatial transcriptomics analyses revealed that SPP1^+^ macrophages and cancer-associated fibroblasts (CAFs) co-localize at the tumor margin, forming an immunosuppressive “tumor immune barrier (TIB)” structure that restricts T-cell infiltration.^25^ Similarly, in pancreatic cancer, multi-omics integration identified a DC-Th-CTL immune niche that could predict responses to chemoradiotherapy.^19^ In bladder cancer, CDH12^+^ epithelial cells were found to engage in spatial interactions with exhausted CD8^+^ T cells via PD-L1/PD-L2 signaling.^26^ Moreover, in glioblastoma, imaging mass cytometry combined with scRNA-seq uncovered spatially distinct myeloid cell subsets associated with long-term patient survival.^27^

Despite significant advances in multi-omics integration, several technical limitations remain. Early spatial transcriptomics platforms, such as 10X Visium, are constrained by their large spot diameter (>55 μm), which hampers the resolution of fine sub-niche structures and obscures localized interactions such as CD40-CD40L mediated B-T cell activation.^28,29^ Although emerging technologies have achieved higher spatial resolution (<10 μm, even reaching subcellular levels, as in MERFISH and Xenium)^30–33^, this often comes at the cost of limited gene detection throughput (<1000 genes), thereby restricting the systematic characterization of regulatory networks within functional niches.^34^ Moreover, a unified framework for multimodal data integration is still lacking. Horizontal integration can suffer from batch-dependent overcorrection, while vertical integration is often constrained by ambiguous feature correspondence across different omic layers. These challenges collectively underscore the need for improved analytical strategies that can balance spatial resolution, molecular coverage, and cross-modal consistency in decoding tissue microenvironments.

Therefore, exploring the heterogeneity of microenvironmental functional niches remains challenged by three major limitations: (1) the unresolved trade-off between spatial resolution and molecular or imaging throughput; (2) the lack of unified and standardized methodologies for multi-omics data integration; and (3) the absence of effective approaches to dynamically trace functional niche remodeling across temporal or therapeutic contexts. To address these challenges, therefore, we developed NICHE, a multimodal integrative strategy that combines single-cell transcriptomics, imaging-based spatial proteomics (such as CmTSA and PhenoCycler-Fusion), and artificial intelligence-driven analytical modeling. This framework enables the construction of high-resolution spatial maps of functional niches and allows the spatial localization of transcriptionally defined cellular subpopulations by integrating spatial interaction features with L-R signaling information at the transcriptomic level. Through this approach, the molecular mechanisms underlying niche organization can be systematically deciphered, and the potential regulatory networks can be reconstructed. Overall, this approach provides a robust avenue for elucidating pathophysiological processes driven by functional niche dynamics, such as immune evasion and therapy resistance, and offers a promising strategy for identifying clinically translatable biomarkers, predicting therapeutic responses, and monitoring disease progression.

## RESULTS

### Construction of single-cell spatial maps and multicellular niche maps of human tonsil based on high-plex CmTSA

Mapping the spatial distribution and dynamic cellular interactions of immune-activating and -suppressive structures within the TiME is crucial for understanding how immune cells exert anti-tumor functions and how tumors evade them. Human tonsil tissue, with its well-defined architecture and diverse immune populations,^35,36^ serves as an ideal model to benchmark the feasibility and robustness of the NICHE analytical strategy. To construct a spatially-resolved cell atlas, we inferred the major immune and stromal populations from a public tonsil scRNA-seq dataset containing 10 samples (**Fig. 1a**). We subsequently developed a 16-marker high-plex panel, which was applied to tonsil FFPE tissues using CmTSA staining (**Fig. 1b, 1c**). Using the high-plex tonsil images, we first employed the deep learning model StarDist-2D for cell segmentation. We then annotated cell types based on biomarker expression patterns and intensity thresholds. This pipeline enabled the digitalization of all major cell types along with their precise spatial coordinates within the tonsil microenvironment (**Fig 1d, e**).

**Fig. 1.**
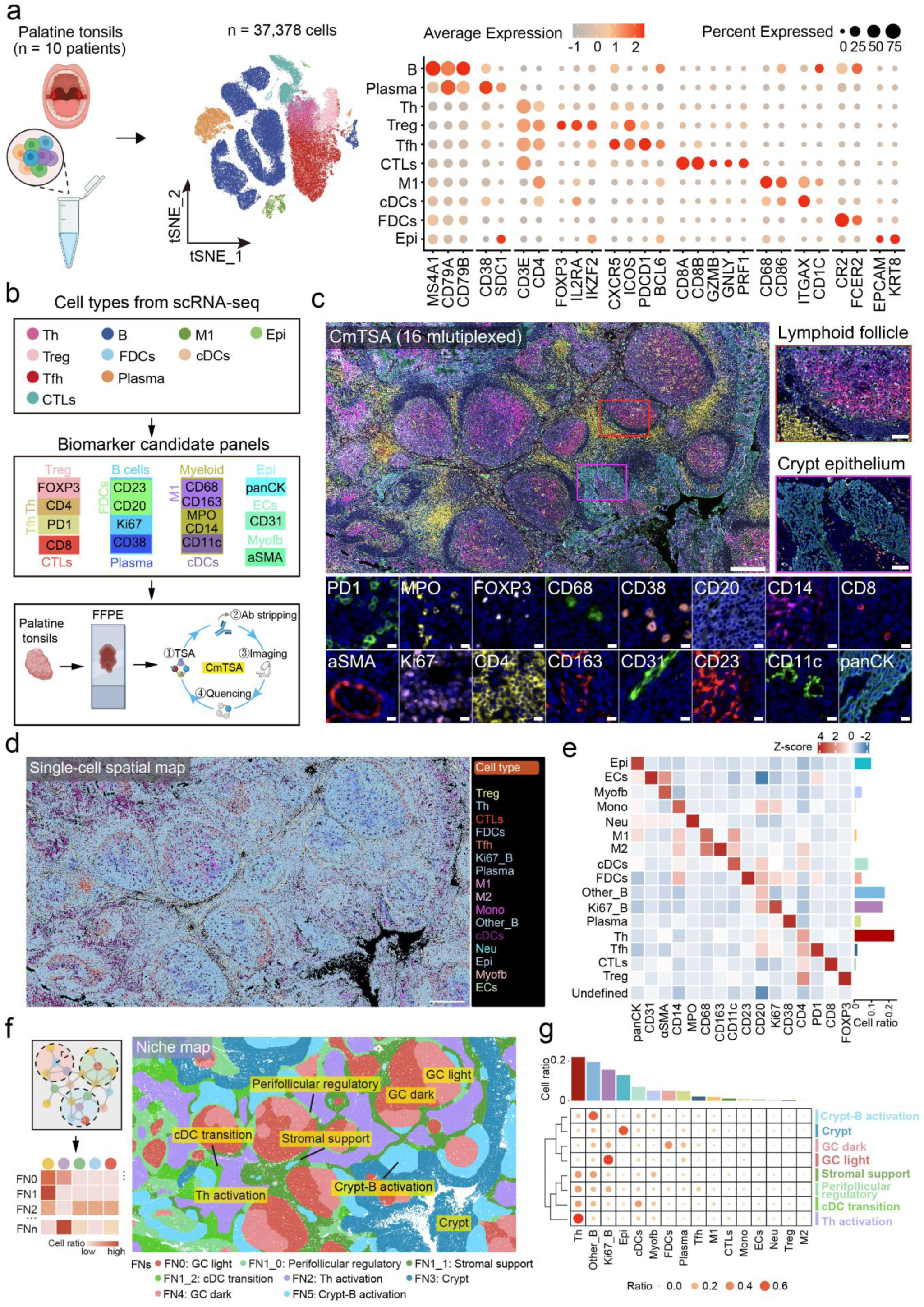
Construction of a single-cell spatial map and FN prediction of human tonsil FFPE tissue using CmTSA. **a** The left panel displays the single-cell sequencing dataset of 10 palatine tonsil samples from public data. The middle panel shows a tSNE dimensionality reduction plot colored by different cell types, and the right panel presents a bubble plot displaying the expression of representative marker genes. Color intensity represents expression level, with red indicating high expression and lighter colors indicating low expression. Bubble size reflects the proportion of cells expressing the gene within each cell type. **b** Antibody panels were designed based on the cell types identified from scRNA-seq data, and multiplex immunofluorescence staining was performed on human tonsil FFPE samples using the CmTSA technique. **c** Representative multiplexed CmTSA stained image of a human tonsil FFPE section showing 16 markers in 17 fluorescent channels. The scale bar for the main image is 500 μm, for the two magnified boxed regions is 100 μm, and for the marker display panels is 10 μm. **d** Definition of cell types in human tonsil tissue based on staining results; scale bar, 500 μm. **e** The heatmap illustrates the expression relationships between different cell types and various protein markers. The Z-score represents the relative expression level of each protein marker across different cell types, with color intensity indicating the magnitude of expression. **f** The left panel illustrates the schematic principle of FN prediction, where each cell is treated as a node to construct a network within a radius r, and the proportion of different cell types in each neighborhood is calculated as clustering features. The right panel shows the predicted spatial FNs of the human tonsil, with different colors representing distinct FNs. **g** The composition of FNs in the human tonsil is shown, where the size and color intensity of each bubble represent the proportion of a given cell type within each FN. Larger and darker bubbles indicate a higher cellular proportion, whereas smaller and lighter bubbles indicate a lower proportion.

Based on the digital single-cell map, we constructed spatial cell networks using a radius-based nearest neighbor (RNN) graph. Cells were grouped into FNs by k-means clustering of their local neighborhood composition. Initial clustering divided the tissue into 6 FNs, one of which (FN1) exhibited marked internal heterogeneity and was further resolved into three sub-niches, yielding a total of 8 spatially and functionally distinct FNs. These computationally derived FNs aligned closely with classical tonsil histology. FN0 (GC light) and FN4 (GC dark) corresponded to the dark and light zones of the germinal center, enriched with Ki67_B/Plasma and FDCs/Other_B, respectively. FN3 (Crypt) matched the crypt epithelium, dominated by epithelial cells (Epi) and cDCs. FN2 (Th activation) represented an extrafollicular Th activation zone, while FN5 (Crypt-B activation) was identified as a crypt-B activation zone. The sub-niches of FN1 delineated transitional regions: FN1_0 (Perifollicular regulatory) as a perifollicular regulatory zone, FN1_1 (Stromal support) as an extrafollicular stromal support zone containing Th, Other_B, and myofibroblasts (Myofb), and FN1_2 (cDC transition) as a cDC transition zone extending toward the subcryptal area (**Fig 1f, g**).

### Defining Spatial Subpopulations through Cell-Cell Interactions within FNs

Various cells in the TiME interact with different cell types in a regular and structured manner, forming specific immune activities that give rise to distinct immune FNs. To gain a deeper understanding of the functions of different FNs within the TiME, we aim to combine single-cell transcriptomic data with functional analysis to extract the communication gene expression profiles of cells within these FNs. In fact, the interacting cell types and the intensity of their interactions vary across different FNs. By integrating single-cell transcriptomic data with CellChat, we can align the true spatial interaction patterns and intensities of cells with the L-R pairs involved in these interactions. This approach allows us to effectively extract spatial subpopulations of specific immune cells within the various FNs. Furthermore, it enables us to explore the functions of different cells within these FNs, as well as the underlying molecular networks that govern their activity and execution.

Local cellular networks were constructed within a 15 μm radius, and interaction strength was quantified using a permutation test, represented as Z-score or log1p(Z-score) to indicate statistical deviation from random association. This approach revealed distinct spatial interaction architectures across FNs. In the interaction heatmap, color intensity reflects the degree of cellular proximity or segregation, while directional arrows suggest asymmetric intercellular relationships, indicative of a potential “encirclement effect” in certain niches. Furthermore, we reconstructed key cellular network structures within each FN, identifying pivotal hub cell types in local communication (**Fig. 2a**). Using this framework, we quantitatively characterized interaction patterns across the 8 FNs. To ensure robustness, only the top 90% of cell types by abundance in each FN were included in the visualization. The analysis revealed distinct communication signatures across niches: FN0 was dominated by Ki67_B-Plasma interactions, implicating it in B cell proliferation and differentiation, whereas the subniches of FN1 exhibited clear functional divergence. FN1_0 showed interactions among Ki67_B, Tfh, and FDCs, suggesting a transitional zone; FN1_1 was characterized by strong Other_B-Myofb interactions, indicative of stromal support and immune regulation; and FN1_2 displayed predominant cDCs-Epi-M1 interactions, supporting a role in antigen presentation. In FN2, interactions were mainly between Th and CTLs, identifying it as a T cell activation core, while FN3 displayed intensive Epi-cDCs-M1 interactions, consistent with the crypt zone. FN4 was dominated by communication among FDCs, Ki67_B, Plasma, and Other_B, reflecting B cell affinity maturation and antibody production. Finally, FN5, characterized by Other_B interactions with Myofb and Ki67_B, was associated with B cell migration and activation near the crypt epithelium (**Fig. 2b**). Overall, the cell-cell interaction profiles align closely with the classical functional architecture of human tonsil, underscoring the utility of our approach in deciphering niche-specific communication patterns.

**Fig. 2.**
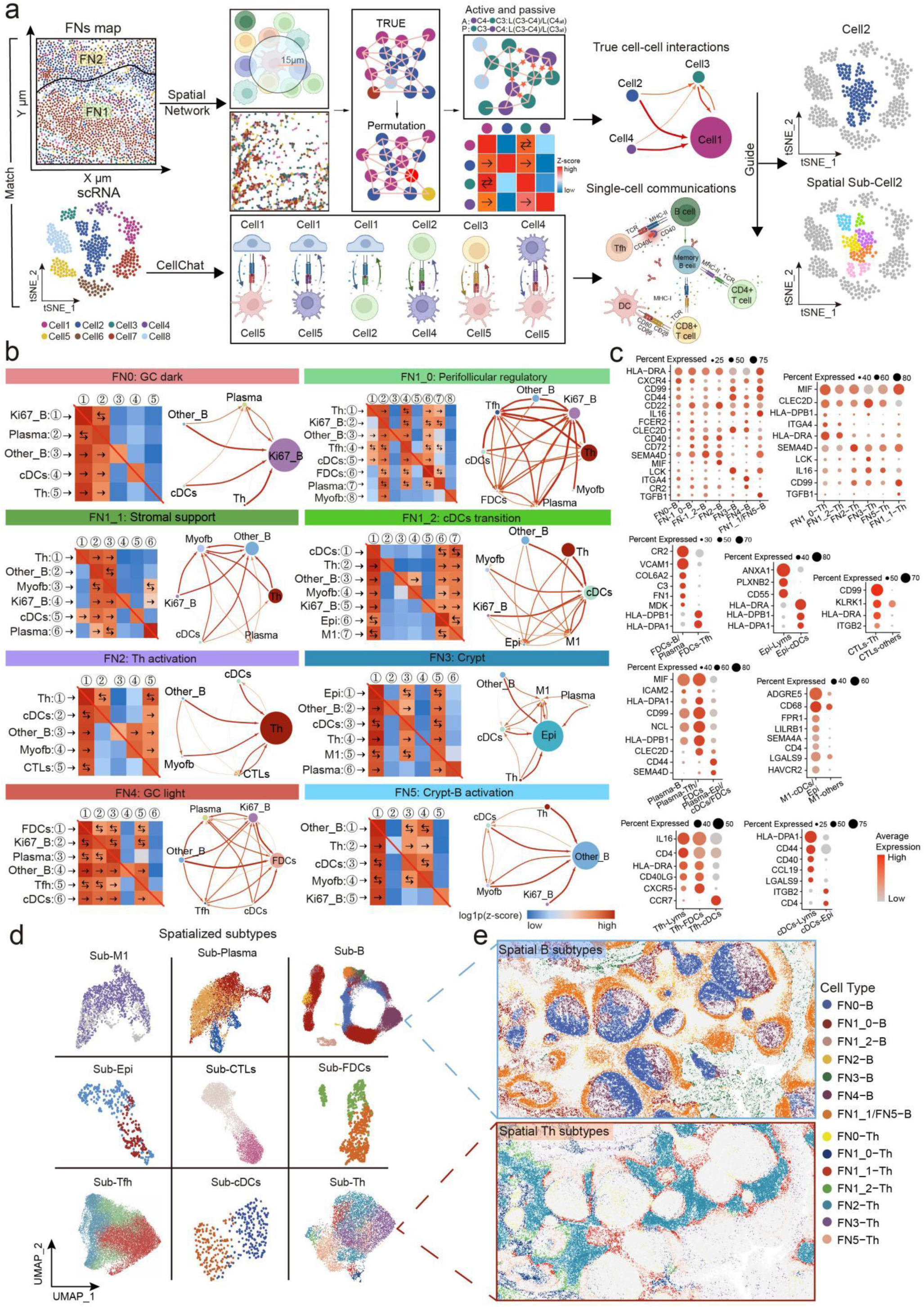
NICHE framework for spatial interactions and mapping of single-cell transcriptomic subpopulations. **a** Schematic illustration of the integration of spatial proteomics and single-cell transcriptomics. The upper panel shows the quantification of cell-cell interactions within FNs by constructing spatial networks. Real and permuted interaction values were compared using permutation tests to obtain Z-scores, which were visualized as heatmaps to represent interaction strength: red indicates stronger cell-cell interactions, while blue represents greater spatial separation. Arrows denote the directionality of interactions, reflecting the spatial dominance of specific cell types. The heatmap information can also be transformed into a network structure to intuitively visualize intercellular interactions within each FN. The lower panel depicts the cell-cell communication network inferred from scRNA-seq data using CellChat, integrating spatial interaction patterns with L-R molecular communications to achieve spatial localization of transcriptional subpopulations within FNs **b** The directional spatial interactions between different cell types within the 8 FNs are shown. The left panel displays a Z-score heatmap, with color intensity representing interaction strength: red indicates a positive interaction trend, while blue indicates no significant interaction or avoidance. Arrows denote the direction of interactions. The right panel shows the corresponding network diagram, where node size reflects the relative proportion of each cell type within the FNs. **c** The bubble plot shows the expression patterns of CellChat communication genes across different cell spatial subpopulations. Subpopulations are located within specific FNs (e.g., FN0-B, FN1-B) or based on cell-cell interactions (e.g., CTLs-Th, CTLs-others). Bubble size represents the proportion of cells expressing the gene, and color intensity reflects the expression level, with red indicating high expression and gray indicating low expression. **d** The UMAP plot shows the distribution of B, Th, FDCs, Epi, CTLs, Tfh, Plasma, M1, and cDCs subpopulations extracted from different FNs in human tonsils. Different colors represent distinct cell subpopulations. **e** The spatial distribution of B and Th subpopulations across different FNs in the human tonsil, corresponding to the transcriptomic subpopulations in (d). Distinct colors represent different cell subpopulations.

To investigate whether the same cell type exhibits functional specialization across different spatial contexts, we visualized the distribution of B and Th cells throughout the tonsil tissue. These cells displayed distinct spatial localization patterns across FNs, suggesting that they may adopt niche-specific functional identities (**Fig. 2e)**. To molecularly decode such spatial specialization, we extracted representative cell-cell interaction patterns from each FN and used them as spatial features to guide the identification of transcriptomic subclusters. Key L-R pairs inferred from CellChat were incorporated as marker genes to align spatial interaction contexts with single-cell transcriptomic data. This integration enabled us to identify transcriptionally distinct subclusters of 9 major cell types including B, Plasma, Tfh, FDCs, cDCs, Epi, M1, Th, and CTLs across the eight FNs, each associated with specific spatial interaction signatures (**Fig. 2c, d**). By establishing a mapping between spatial interaction features and transcriptomic subclusters, we achieved multi-omic alignment of cellular states across modalities, providing a foundation for elucidating the molecular mechanisms underlying the functional organization of FNs.

### High-Resolution Molecular Profiling Defines the Tonsil Crypt as a Dual-Function Immune Niche

To further evaluate the utility of NICHE in deciphering FNs in tissue microenvironments, we applied it to molecularly deconstruct the crypt zone (FN3) of human tonsil—a key site in mucosal immunity (**Fig. 3a**). Although Epi constituted the most abundant population in FN3, spatial network analysis revealed that Epi, cDCs, and M1 macrophages all exhibited high network centrality, indicating their roles as hub cells in local interaction circuits. We subsequently identified transcriptomic subpopulations of these hub cells within FN3. Gene Ontology (GO) enrichment analysis of their gene expression signatures classified the functional profile of the crypt zone into six major categories: positive immune regulation, antigen presentation, macrophage and autophagy-related processes, tissue repair, negative immune regulation, and apoptotic signaling. This functional spectrum reflects the multifaceted role of the crypt in coordinating immunity and maintaining tissue homeostasis (**Fig. 3b**). Differential gene expression analysis of Epi, cDCs, and M1 subpopulations further revealed upregulated expression of immune-related genes including *S100A7/8/9, CXCL10, LYZ, CXCL6, CCR7, IL1B, TLR2, IL36RN, IL36G, EGFR, CXCL17, CTSD, HLA-DRA, CCL19,* and *TLR4*, which are implicated in antimicrobial defense, chemotaxis, antigen presentation, and tissue remodeling. Additionally, we detected expression of immunosuppressive or escape factors such as *TGFB2, IDO1, AHR,* and *LGALS3*, suggesting that the crypt zone not only initiates immune responses but also engages active regulatory mechanisms to maintain local immune homeostasis (**Fig. 3c**). Multiplex immunofluorescence staining further validated the expression of these key genes within crypt zone at the protein level (**Fig. 3d**).

**Fig. 3.**
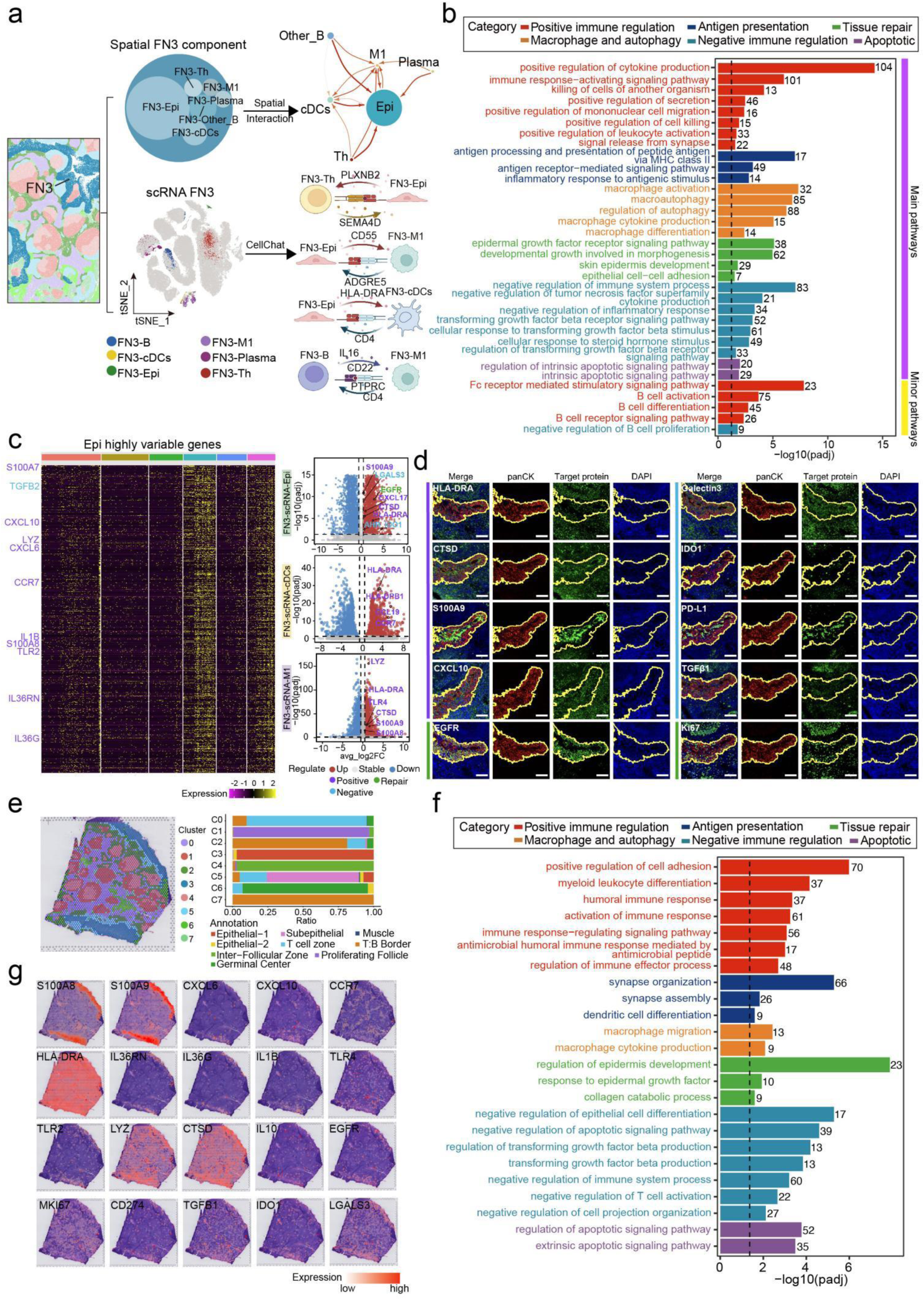
Analysis of the immune features and molecular mechanisms of the human tonsil crypt (FN3) **a** The crypt data includes both spatial subpopulations and transcriptomic subpopulations. The upper panel shows the spatial subpopulation information, while the lower panel presents the UMAP dimensionality reduction of transcriptomic subpopulations, with cells colored according to subpopulation type. **b** Bar plot showing the enrichment of major and minor pathways in the FN3 crypt FN. The bars represent the significance level of each pathway (−log10(padj)), with different colors indicating distinct pathway categories. Purple bars on the right denote main pathways, while yellow bars represent minor pathways. **c** Left: Heatmap of highly variable gene expression in the FN3-Epi cluster, with yellow for high expression and purple for low expression. Right: Volcano plots of differentially expressed genes in the Epi, cDC, and M1 subpopulations in FN3, with log2 fold change (x-axis) and expression significance (−log10(padj), y-axis). Red and blue dots indicate upregulated and downregulated genes, respectively, while gray dots represent non-significant genes. Genes are colored by function: purple for positive regulation, green for repair-related, and blue for negative regulation. **d** CmTSA staining revealed the expression of multiple proteins in the crypt zone (panCK marking the crypt zone), including markers of immune activation (HLA-DR, CTSD, S100A9, CXCL10), tissue repair (EGFR, Ki67), and immune negative regulation (Galectin-3, IDO1, PD-L1, TGFβ1). Scale bar, 100 μm. **e** The left panel shows the SpatialDimPlot of human tonsil spatial transcriptomic data, where different colors represent distinct cell clusters (Cluster 0–7). The right panel presents the proportion of various cell types within each cluster, with colors corresponding to different cell types. **f** The plot shows the pathway enrichment analysis of the C3 crypt zone. Bar lengths represent the significance level of each pathway (−log10(padj)), and different colors indicate distinct pathway categories. **g** The C3 crypt zone is shown first, with each panel illustrating the spatial expression patterns of different genes within FN3. Color intensity represents gene expression levels, where orange-red indicates high expression and white indicates low expression.

To further Orthogonal validate these findings, we compared the NICHE results with spatial transcriptomic (ST) data of human tonsil. A high degree of concordance was observed at both the niche and molecular levels. The crypt zone (C3) identified by ST closely matched FN3 from NICHE in terms of spatial localization and cellular composition (**Fig. 3e**). Functionally, both methods revealed similar pathways, including positive immune regulation, antigen presentation, and negative immune regulation (**Fig. 3f**). In addition, key genes from FN3 derived from NICHE, such as HLA-DRA, S100A9, CXCL10, and CTSD (immune activation), as well as LGALS3, IDO1, and TGFB1 (immune suppression), were expressed to varying degrees in the C3 region, further validating the dual role of the crypt in immune activation and regulation (**Fig. 3d, g**). Collectively, these results firmly establish the tonsil crypt not as a passive barrier, but as a dynamic immune hub equipped with a balanced “accelerator-and-brake” mechanism. This precise co-expression of activating and inhibitory molecules enables robust defense while preventing immunopathology, the disruption of which may underlie mucosal immune dysregulation. It should be noted that while ST data provided valuable population-level validation, the NICHE framework operates at single-cell resolution, offering deeper and more robust molecular profiling and enabling higher-resolution analysis of cellular interactions.

### Tracing the CD8_T cytotoxic niche within the TiME of NSCLC before and after neoadjuvant immunotherapy

The NICHE framework enables robust profiling of FNs crosstalk at molecular and cellular levels, providing a powerful approach to monitor key FNs in dynamic settings. Here, we applied NICHE to track the CD8_T cytotoxic niches in NSCLC patients undergoing neoadjuvant immune checkpoint blockade—a niche critically linked to divergent pathological responses, including pathologic complete response (pCR), major pathologic response (MPR), and partial pathologic response (pPR). We included FFPE samples from 16 NSCLC patients who underwent neoadjuvant immunotherapy, with 4 samples used for the discovery group and 12 samples used for the validation group (**Fig. 4a**). Additionally, we utilized a publicly available NSCLC scRNA-seq dataset, which encompassed both pre- and post-treatment samples representing the pPR, MPR, and pCR states. The scRNA-seq analysis revealed the major components of the TiME, including immune, myeloid, mesenchymal, and epithelial (both normal and malignant) cell populations (**Fig. 4b**). Based on this, we designed a 19-biomarker CmTSA panel and applied it to 4 NSCLC FFPE samples from the discovery group: 1 pre-treatment and 3 post-treatment (pPR, MPR, pCR)(**Fig. 4d**). Pathologist-annotated regions of response on serial H&E-stained sections guided our subsequent spatial analysis (**Fig. 4c**). We generated single-cell spatial maps and quantified cell populations across all samples (P1-Pre: 194,472; P2-pPR: 58,668; P3-MPR: 147,129; P4-pCR: 294,514)(**Fig. 4e**).

**Fig. 4.**
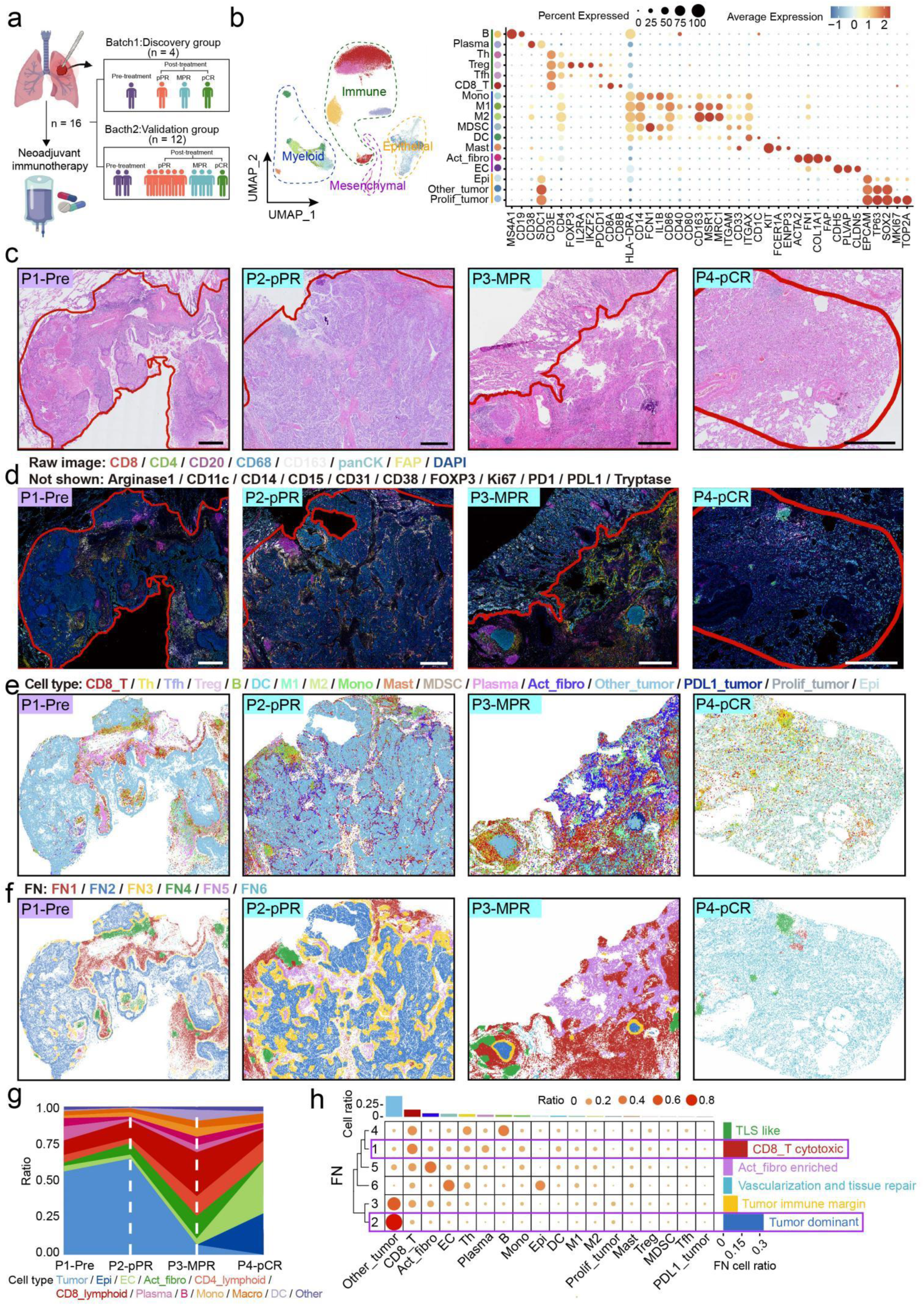
Pre- and post-neoadjuvant immunotherapy TiME FNs mapping and analysis in NSCLC. **a** The inclusion of clinical FFPE samples from NSCLC in neoadjuvant immunotherapy, including the pre-treatment group and post-treatment subgroups with different therapeutic responses: pPR, MPR, and pCR. **b** UMAP plot of scRNA-seq cell annotations from public NSCLC datasets, with colors representing different cell types. The right bubble plot shows marker gene expression patterns, with bubble size indicating the proportion of expressing cells and color intensity reflecting expression levels, red for high expression, blue for low expression. **c** H&E stained image of a lung cancer FFPE section. The red outline indicates the tumor response region annotated by the pathologist. Scale bar, 1 mm. **d** Representative image of CmTSA 18 markers staining performed on an H&E adjacent serial section of a lung cancer FFPE sample. The red outline indicates the tumor response region annotated by the pathologist. Scale bar, 1 mm. **e** Spatial mapping of cell types within the tumor response region, with different colors representing distinct cell populations. **f** Network constructed within a 100 μm radius and projection map of FNs clustered based on the neighborhood composition of each central cell. **g** Relative proportions of major cell categories in pre-treatment and post-treatment samples with different therapeutic responses, with different colors representing distinct cell types. **h** Cell composition of different FNs, with bubble size and color intensity representing the proportion of each cell type.

Cellular composition analysis showed tumor cells predominating pre-treatment, with a progressive decline post-therapy accompanied by increased immune and stromal components (**Fig. 4g**). Spatial analysis identified 6 recurrent FNs across samples (**Fig. 4f**), including a CD8_T cytotoxic FN, a tertiary lymphoid structure (TLS) like FN, a tumor immune margin FN, a tumor dominant FN, an activated fibroblast enriched (Act_fibro enriched) FN, and a vascularization and tissue repair FN (Figure 4H). The CD8_T cytotoxic FN, enriched in CD8_T, Th, B, Plasma, and dendritic cell (DC)—was identified as the principal niche responsible for tumor cell killing. It localized adjacent to tumor mass regions (tumor dominant and tumor immune margin FNs) and exhibited a dynamic abundance across response groups: intermediate in pre-treatment and pPR, highest in MPR, and nearly absent in pCR, reflecting its evolving role during treatment. (**Fig. 4e-h**).

### Profiling of the Dynamic Cellular Remodeling of CD8_T cytotoxic niche across Treatment Responses

Using the NICHE framework, we characterized the spatial and cellular evolution of CD8_T cytotoxic FN across pre- and post-treatment response states (pPR, MPR, pCR)(**Fig. 5a**). Comparative analysis revealed a marked dynamic progression in network architecture. In pretreatment samples (Pre), the cellular interaction network was highly complex, involving multiple cell type, including Plasma, CD8_T, Th, Monocyte (Mono), M1, and M2 macrophages, with Plasma as the dominant participants. Following neoadjuvant immunotherapy, the pPR sample exhibited a shifted landscape: CD8_T expanded and became the dominant population, engaging not only with tumor cells but also with monocytes and an increased population of activated fibroblasts (Act_fibro) in the pPR state. These fibroblasts displayed elevated network centrality, interacting strongly with Th, Mono and CD8_T, indicating active stromal-immune crosstalk. By the MPR state, the interaction network simplified considerably, dominated by CD8_T, Th, and DC, with Act_fibro and Mono assuming secondary roles. In pCR state, the network further condensed into a compact core centered on CD8_T-Th-DC interactions. It is noteworthy that in both the pCR CD8_T cytotoxic FN (FN1) and vascularization and tissue repair FN (FN6), CD8_T no longer forms dense multicellular clusters (**Fig. 5b-d and S3**). We found that CD8_T and Act_fibro underwent the most significant dynamic changes.

**Fig. 5.**
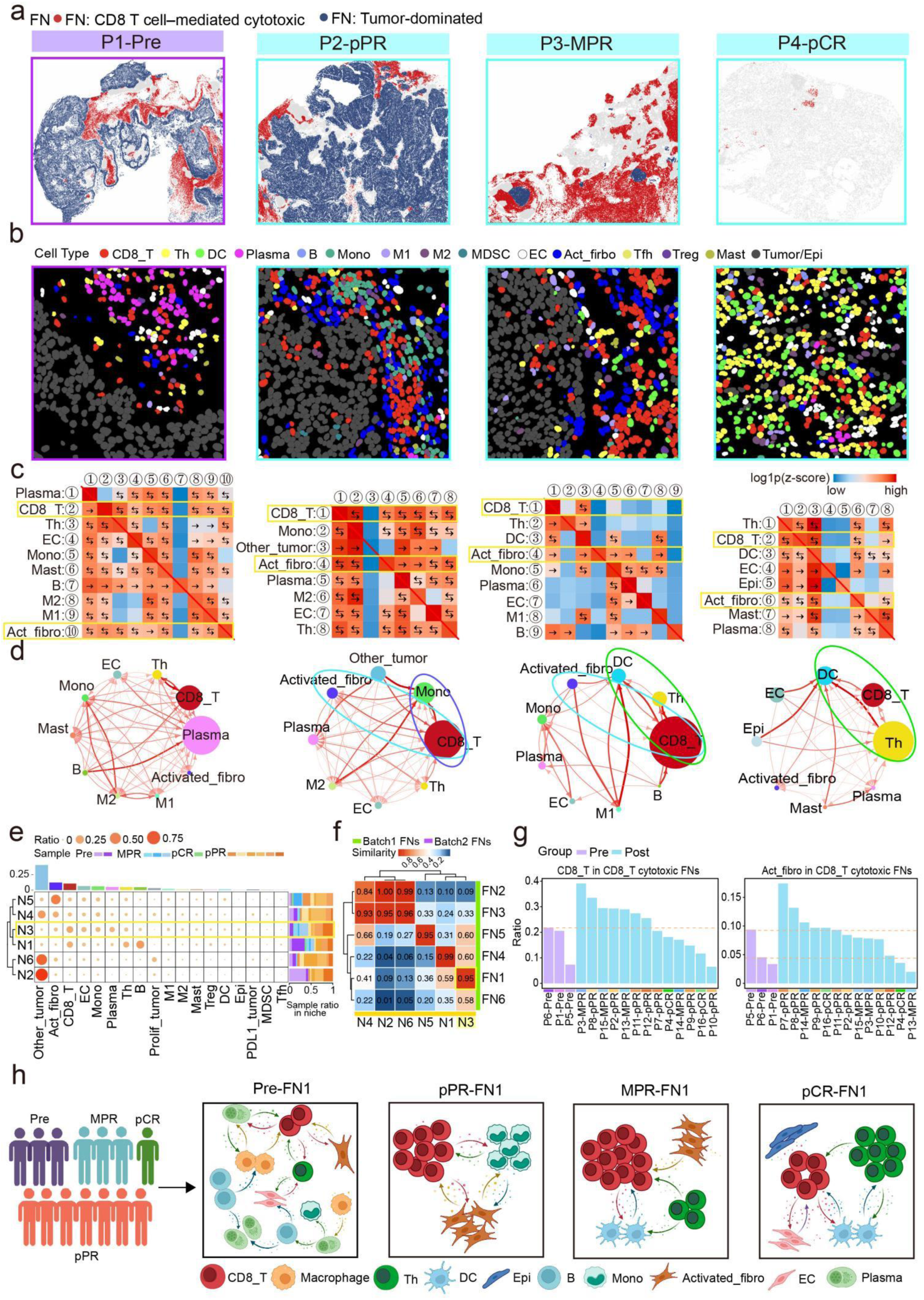
Spatial architecture of the CD8_T cytotoxic FN (FN1) **a** Spatial projection map of the CD8_T cytotoxic FN (red) and tumor-associated FN (dark blue) across different samples. **b** The representative in situ cell images of the FN1 region for each sample are shown. **c** Heatmap of directional spatial interactions between different cell types within the CD8_T cytotoxic FN across samples, showing log-transformed Z-scores. Color intensity represents interaction strength, with red for positive interactions and blue for no significant interaction or avoidance. Arrows indicate the direction of interactions.. **d** Cell interaction network diagrams, where node size represents the proportion of each cell type within the CD8_T cytotoxic FN. **e** Cell composition of different functional niches(Ns), with bubble size and color intensity representing the proportion of each cell type. The bar chart on the right represents the distribution of functional niches across patients. **f** The cosine similarity of 6 functional niches between Batch 1 discovery group and Batch 2 validation group, with red indicating higher similarity. **g** The ratio of CD8_T and Act_fibro within the CD8_T cytotoxic FNs across all patients are shown. **h** Schematic diagram of cell interactions within the CD8_T cytotoxic FNs across different groups.

Therefore, we performed a validation analysis of CD8_T and Act_fibro significance in the 12 patient samples from the validation group (including 2 Pre, 6 pPR, 3 MPR, and 1 pCR). The samples from the validation group were collectively divided into 6 functional niches (Ns), defined as N1-N6 (**Fig. 5e and S4**). To correspond with the functional niches (FN1-FN6) of the discovery group, we calculated the cosine similarity of the 6 functional niches in both groups and found that N3 in the validation group corresponds to the previously defined CD8_T cytotoxic FN (FN1) (**Fig. 5f**). We then calculated the ratio of CD8_T cytotoxic and Act_fibro in the 16 samples from the discovery and validation groups. Compared to the pre-treatment samples, post-treatment samples showed a synchronized increase in the ratio of CD8_T and Act_fibro (**Fig. 5g**). Overall, spatial interaction patterns transitioned from broad complexity to a focused, specialized structure. The dynamic changes in CD8_T and Act_fibro were validated in the expanded cohort, underscoring their pivotal roles in mediating cytotoxicity and stromal regulation during therapy (**Fig. 5h**).

### CD8_T-Act_fibro negative feedback interaction shapes the dual-state cytotoxic FN after therapy

To molecularly decode these spatial changes, we integrated key cellular interaction pairs within each FN with the L-R networks inferred by CellChat. This enabled us to identify critical signaling molecules, which were used as feature genes to map the spatial subclusters onto the scRNA-seq data. Through this approach, we obtained various transcriptomic cell subpopulations corresponding to the CD8_T cytotoxic FNs in different samples (**Fig 6a, b and S5, 6**). Based on GO pathway enrichment analysis of differentially expressed genes from key spatial subpopulations within the cytotoxic FNs, including CD8_T, DC, Mono, Act_fibro, M2, Th, Plasma, and EC, we found that this FN was significantly enriched in immune related pathways, such as CD8_T anti-tumor response, T cell commitment, and myeloid immune activation. Concurrently, pathways associated with tissue repair and negative immune regulation were also observed (**Fig. 6c**). These findings indicate that, following therapy, the cytotoxic FNs exhibits a complex dual state characterized by both enhanced antitumor immunity and concomitant activation of repair and immunosuppressive signals. Building on this foundation, we further focused on the 2 cell types that exhibited the most significant changes pre- and post-treatment, CD8_T and Act_fibro. Differential gene expression analysis between pre- and post-treatment samples revealed that in CD8_T, cytotoxic effector molecules such as *GZMA, GZMB, PRF1, GZMH, CTSW*, and *EOMES* were markedly upregulated after treatment, indicating enhanced cytotoxic activity. Meanwhile, exhaustion markers including *LAG3, TIGIT, PDCD1, HAVCR2*, and *ADGRG1* were also elevated, suggesting that CD8_T exhibited a coexisting state of activation and exhaustion. In Act_fibro, genes associated with tissue repair and extracellular matrix (ECM) remodeling, such as *COL15A1, TNC, COL6A1, TAGLN2, POSTN, TGM2, LTBP2, ADAMTS2*, and *TGFBI* were significantly upregulated post-treatment, indicating enhanced matrix remodeling and tissue repair activity. At the same time, the increased expression of inhibitory factors such as *HIF1A, CXCL12*, and *CXCL16* suggests that Act_fibro may also exhibit suppressive functions during the repair process (**Fig. 6d**). By further applying CellChat to infer signaling networks mediating the interactions between CD8_T and Act_fibro within the post-treatment cytotoxic FNs, we identified the top-ranked information flow pathways. These included classical immune regulation signals such as *MHC-I, CCL, LCK, MHC-II, CXCL, IFNG-II*, and *CD226*, as well as ECM and matrix remodeling associated pathways including *COLLAGEN, ITGB2, LAMININ, FN1,* and *PERIOSTIN*. In addition, several immunosuppressive or exhaustion-related signals, such as *NECTIN, TIGIT, SEMA3/4,* and *TGFb*, were also enriched, indicating a complex signaling landscape that integrates immune regulation, matrix remodeling, and immune resistance within the cytotoxic FNs (**Fig. 6e**). We inferred the L-R interaction pairs between CD8_T and Act_fibro, as well as between Act_fibro and CD8_T, and proposed a bidirectional CD8_T-Act_fibro interaction model based on CellChat analysis. On one hand, CD8_T cells may activate Act_fibro through signaling pathways such as IFNG-(IFNGR1+IFNGR2), SEMA4D-PLXNB2, and GZMA-PARD3, potentially promoting ECM remodeling, cell polarity regulation, and tissue repair. On the other hand, Act_fibro might transmit inhibitory feedback through various mechanisms, including the NECTIN2-TIGIT immune checkpoint axis, the CXCL12-CXCR4 chemotactic barrier, and several ECM-receptor interactions, such as FN1-CD44, COL6A3-CD44, and COL6A3-(ITGA1+ITGB1), collectively possibly limiting CD8_T infiltration and effector activity (**Fig. 6f**). Furthermore, through observations from CmTSA staining results, we found that fibrous encapsulation at the tumor margins of residual tumors hindered CD8_T infiltration. In the inflammatory environment, fibroblast activation may jointly restrict CD8_T cell infiltration at both the molecular and physical levels, which is consistent with our findings inferred from scRNA-seq data analysis (**Fig. 6g**).

**Fig. 6.**
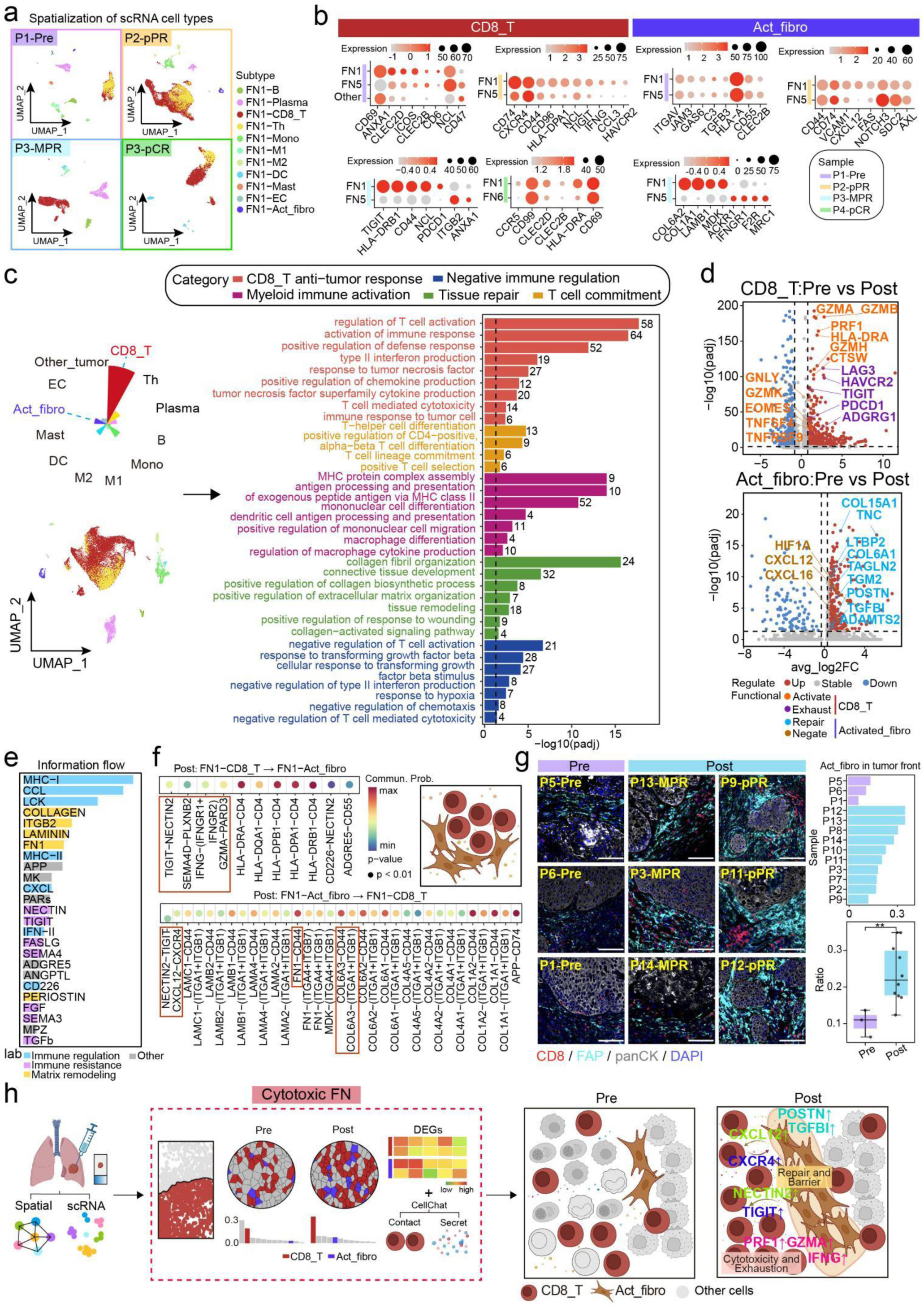
Exploration of molecular mechanisms in CD8_T cytotoxic FN (FN1) **a** UMAP plot of scRNA-seq data from FN1, with colors representing different cell subpopulations. **b** CellChat gene expression in CD8_T and Act_fibro across different FNs, with red indicating higher expression and larger bubbles representing higher expression proportions. **c** Spatial cell composition of post-treatment FN1 and its alignment with spatially localized single-cell transcriptomic subpopulations, along with a summary of GO enrichment analysis for key cell types. The color of the bar chart represents different biological functions. **d** Volcano plots illustrate the upregulated and downregulated genes in the spatial subclusters of CD8_T (top) and Act_fibro (bottom) before and after treatment. The highlighted key genes represent typical cellular functional states, including Activate, Exhaust, Repair, and Negative. **e** Information flow analysis using CellChat illustrates the relative strength of major signaling pathways between CD8_T and Act_fibro cells in post-treatment samples, where the height of each bar represents the magnitude of pathway information flow. **f** L-R interactions between CD8_T and Act_fibro cells within the FN1 of post-treatment samples, where color intensity represents the strength of communication probability and circle size indicates significance (p < 0.01). Red boxes highlight key signaling axes involved in immune regulation. **g** On the left, representative images of Act_fibro in the tumor front are shown for pre-and post-treatment samples, with a scale bar of 100 μm. On the right, the ratio of Act_fibro in the tumor front across all samples is quantified, with statistical significance of 0.036. **h** Summary schematic of the NICHE framework applied to lung cancer, providing a multidimensional analysis of the CD8_T-Act_fibro negative feedback signaling.

Taken together, these findings suggest the presence of a potential negative feedback loop between CD8_T and Act_fibro within the cytotoxic FN, where CD8_T cells may activate stromal responses, while Act_fibro may exerts counter-regulatory suppression through physical barriers and chemotactic factors. This mechanism may partially explain why residual tumor cells persist after immunotherapy. We further examined the expression of the aforementioned differentially expressed genes (DEGs) and L-R pairs of CD8_T and Act_fibro in post-treatment scRNA-seq data. In CD8_T cells, cytotoxic and communication associated molecules (such as IFNG, GZMA, SEMA4D, CD44, and ITGA1), as well as exhaustion and chemotaxis related factors (such as TIGIT, PDCD1, and CXCR4), exhibited high expression levels. In Act_fibro, receptor and ECM related molecules (such as IFNGR1, PARD3, PLXNB2, COL6A1, FN1, and POSTN), along with immunosuppressive axis components NECTIN2 and CXCL12, also showed high expression levels (**Fig. S7**).

Finally, using the NICHE framework, we illustrated how spatial organization and molecular signaling shape the CD8_T cytotoxic FN and speculated on the bidirectional influence between CD8_T and Act_fibro. This bidirectional influence may establish a dynamic balance between immune activation and stromal suppression, providing a certain degree of theoretical basis for understanding the persistence of residual tumors in NSCLC patients after neoadjuvant immunotherapy (**Fig. 6h**).

## DISCUSSION

This study focused on integrating single-cell transcriptomics with spatial proteomics to investigate cellular subpopulation interactions within the tissue microenvironment, address the challenges of cross-modal data alignment, and highlight the methodological advances achieved through this integration.

Traditional single-cell transcriptomics enables high-resolution profiling of cellular transcriptional states but lacks spatial context. Consequently, whether the identified cellular subpopulations truly exist within the native tissue architecture and how they are spatially distributed remain critical questions. In this work, we introduced a L-R based transcriptomic–spatial concordance validation strategy to ensure that the defined cellular subpopulations possess not only distinct molecular signatures but also verifiable spatial localization. This approach effectively aligns transcriptomic findings with spatial biological reality, thereby enhancing the biological validity of cellular subpopulation classification.

Current mainstream spatial omics platforms (e.g., 10× Visium) remain constrained by spot-level mixed signals, which limit their ability to achieve true single-cell resolution.^37^ This study proposes two complementary directions for improvement. First, on the experimental side, higher-resolution spatial technologies (such as MERFISH^38^ and CosMx^39^ or single-molecule FISH^40,41^ validation can be introduced to verify the spatial localization of key marker genes. Second, on the computational side, deconvolution algorithms (e.g., SpaDecon^42^ and RCTD^43^ can be applied to infer the cellular composition within each spot, thereby establishing a finer correspondence between spatial and scRNA data. Although these methods still face challenges such as limited depth and ambiguous spatial boundaries, their combined use can substantially improve spatial resolution and interpretability.

To address the bottleneck of cross-modal alignment, this study introduces a multi-level integration strategy. Specifically, spatial matrix data were first used to quantify directional cell-cell interactions within each FN. Subsequently, communication networks constructed by CellChat in the scRNA-seq dataset were employed to identify L-R signaling molecules as molecular markers. These markers were then integrated with spatial interaction features to guide the spatial localization of single-cell subpopulations, enabling precise classification of spatially resolved subclusters. This strategy provides a novel and generalizable solution for cross-modal data integration and alignment.

In summary, at the technical level, our analytical framework achieved two major breakthroughs: (1) By leveraging CellChat-based L-R network analysis, we were able to distinguish cell subpopulation-specific interaction signals and uncover the hierarchical communication architecture of the immune microenvironment; (2) Through the integration of transcriptomic, spatial, and functional annotations, we progressively constructed a multi-dimensional evidence chain, enabling a systematic interpretation of the “identity–location–function” relationships of cells within complex tissue ecosystems. This workflow provides a novel pathway for cross-modal data alignment and establishes a methodological foundation for future studies of the tissue microenvironment. At the biological level, our study revealed differences in immune dynamics across patients with different therapeutic outcomes following neoadjuvant immunotherapy for NSCLC. In pPR patients, the CD8_T cytotoxic FN failed to drive complete remission, likely due to the enhanced CD8_T cytotoxic activity that concurrently activated Act_fibro, which, in turn, reinforced the ECM and secreted immunosuppressive factors, thereby limiting the sustained cytotoxic function of CD8_T. This finding provides important insights into the mechanisms underlying immune response failure and suggests potential therapeutic strategies targeting activated fibroblasts or their downstream signaling pathways, which could help mitigate stromal barriers and immune suppression, thereby improving immunotherapy efficacy. Overall, this integrative strategy provides a conceptual framework for basic immunology and a theoretical basis for advancing precision diagnostics and therapeutic interventions in immune-related diseases.

## MATERIALS AND METHODS

### FFPE tissue section preparation

FFPE tissue sections of human tonsil and NSCLC samples (**supplementary Table S1**) were obtained from West China Hospital, Sichuan University, following approval by the institutional ethics committee.

### H&E and CmTSA staining of histopathological sections

For each paraffin-embedded tissue block, 2 consecutive sections with a thickness of 3.5 μm were prepared. One section was subjected to H&E staining, scanned, and annotated by a pathologist to identify tumor response regions. The adjacent section was processed using the CmTSA staining protocol to obtain single-cell resolution spatial proteomic information from the tissue section.^44^

### Antibody infromation

See supplementary Table S2.

### Nuclear segmentation and identification of positive cells

All image-based analyses were performed using QuPath (v0.5.1) software^45^. The operation and workflow of QuPath followed the official documentation and guidelines available at https://QuPath.readthedocs.io/en/stable/.

### Cell segmentation

A new analysis project was created, and the images to be analyzed were imported into the project. Each channel within the images was assigned a specific name, with the nuclear channel used for cell segmentation designated as “DAPI.” Regions of interest (ROIs) were manually selected for analysis. The pre-trained StarDist^46^ model parameters were configured, and the model was executed to perform segmentation (**Fig. S1a**). Upon completion, nuclear segmentation results were obtained within the defined ROIs.

### Identification of positive cells

Based on the nuclear segmentation results, fluorescence intensity values for each channel were obtained for all cells. Thresholds were then defined according to the fluorescence intensity of each marker to distinguish positive from negative cells, and each threshold was saved as an independent classifier. After threshold calibration for all marker channels, all classifiers were loaded simultaneously to identify positive signals for each cell. Upon completion, all operations were saved at the project level, and per-cell vector data were exported for downstream spatial analyses. The exported data included: Object ID, Classification, Parent, Centroid X μm, and Centroid Y μm. Using the cell vector data exported from QuPath, annotation rules were defined to assign cell identities based on staining quality and cellular lineage features, thereby determining the specific cell type of each segmented cell (**Fig. S1a, b**).

### Acquisition of scRNA-seq data

The scRNA-seq dataset of human tonsil used in this study was obtained from publicly available resources deposited on the Zenodo platform.^47^ The scRNA-seq dataset of human lung cancer (GSE207422) was retrieved from publicly accessible resources hosted by the National Center for Biotechnology Information (NCBI).^48^ During data acquisition, all procedures strictly followed the usage guidelines of the original studies, ensuring the integrity and accuracy of the retrieved data.

### Cell annotation of scRNA-seq data

After removing doublets and cells with high mitochondrial gene expression, data processing was performed using Seurat (v4.3.0.1). The dataset was first normalized, and the top 2,000 highly variable genes were selected for downstream analysis. To correct for batch effects, data integration was carried out using the Harmony algorithm. Subsequently, principal component analysis (PCA) was conducted using the top 20 principal components, followed by UMAP or tSNE dimensionality reduction and cell clustering. Finally, clusters were annotated based on the expression of canonical marker genes reported in the literature.

### Prediction of intercellular communication molecules using CellChat

After obtaining the spatial interaction profiles of FNs, we selected key cell types within each FN as the focus for subcluster identification. Subsequently, CellChat (v1.6.1) was employed under default parameter settings to infer intercellular communication networks based on the Seurat objects. L-R signaling pairs corresponding to major spatial interactions were extracted from the CellChat predicted network, and these communication molecules were used as marker genes to guide the spatial mapping of single-cell transcriptomic subclusters. Through this approach, we identified transcriptomic subpopulations with defined spatial localization within functional niches.

Differential gene expression analysis of spatially resolved single-cell subpopulations Each spatially localized cellular subpopulation was processed using the standard Seurat workflow, and differential gene expression analysis was conducted with the FindMarkers function. To enhance analytical accuracy, the logfc.threshold parameter was set to 0.25 to define the minimum threshold for gene expression changes. Genes were filtered using an adjusted p-value < 0.05 and subpopulation specific avg_log2FC cutoffs, depending on the analytical context, to identify significantly differentially expressed genes. Finally, volcano plots were generated using ggplot2 (v3.5.1), with major differentially expressed genes labeled for visualization.

### GO analysis of scRNA-seq subpopulations

Significantly differentially expressed genes identified in each cellular subpopulation were subjected to enrichment analysis using the enrichGO function in the clusterProfiler (v4.8.3) package. The parameter OrgDb = org.Hs.eg.db was specified to obtain human gene annotation information, and the “BP” (Biological Process) category was selected to perform Gene Ontology based biological process enrichment analysis.

### FNs prediction based on single-cell spatial proteomics data

Building upon the FN prediction principle of the spatial omics analysis tool SOAPy (v0.1.4),^49^ the method was appropriately refined to better accommodate the laboratory’s requirements for large sample sizes and parallel analyses, enabling more efficient functional niche prediction. Specifically, a spatial neighborhood network was constructed for each cell using a 100 μm radius. This relatively large distance scale was chosen to capture macroscopic spatial organization and functional niche patterns across tissue sections containing hundreds of thousands of cells. To ensure network robustness, only neighborhoods containing at least 5 valid cells were retained, while those below this threshold were filtered out. Based on the compositional features of neighboring cells within this radius, k-means clustering (using default parameters) was applied for cell grouping. For both human tonsil and lung cancer samples, the number of clusters was adjusted according to sample characteristics to obtain FN delineations that accurately reflect macroscopic tissue structural organization.

### Cosine similarity based matching between FNs and Ns

After dividing the 12 samples in the batch2 validation group into 6 functional niches, the cellular composition of each niche was obtained. Cosine similarity was then used to compare the 6 functional niches (FNs) identified in the discovery group with the 6 functional niches (Ns) identified in the validation group. Based on these similarity scores, the niche in the validation group most closely resembling the CD8_T cytotoxic functional niche was identified.

### Construction of cell-cell networks within FNs

After extracting the spatial coordinates of cells within each FN, the R packages FNN (v1.1.3.2), Matrix (v1.6-1.1), and igraph (v1.5.1) were used to determine cell-cell neighborhood relationships. For each cell, a local spatial range with a radius of 15 μm was defined as the proximity threshold for potential intercellular interactions, representing the distance within which cell-cell communication is likely to occur. Based on these relationships, a spatial adjacency matrix was constructed at the single-cell level to characterize the potential intercellular interaction network within each FN.

### Quantification of cell-cell interactions within FNs

To investigate the directional interaction strength among different cell types within each FN, we developed an Interaction Tendency Score (ITS) based on previously reported methodologies.^49,50^ Using the established spatial cell network, the interaction score from cell 1 to cell 2 was defined as the ratio of the number of connecting edges between the two cells to the total number of edges of cell 1; conversely, the interaction score from cell 2 to cell 1 was defined as the number of connecting edges between the two cells divided by the total number of edges of cell 2. This metric quantifies the directional intensity of intercellular interactions. To assess the statistical significance of the interaction scores, a permutation test was performed. Specifically, the spatial coordinates of the cells within the network were randomly permuted to generate a set of randomized networks, and this process was repeated 100 times (n = 100). For each permutation, interaction scores were recalculated to construct a null distribution, and the real interaction scores were then compared against this randomized background. Standardization was applied to eliminate random noise, allowing for the identification of statistically significant directional interactions between cell types within each FN. Moreover, since the interactions from cell 1 to cell 2 and cell 2 to cell 1 exhibit inherent directionality, this computational framework conceptually parallels the source-target paradigm used in CellChat, thereby providing a biologically meaningful representation of intercellular signaling directionality. The formula is as follows:

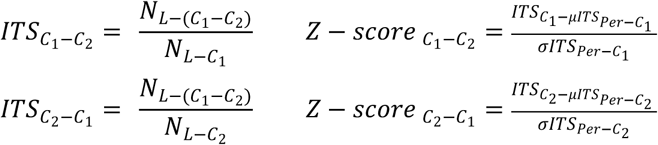

In the equation 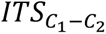 and 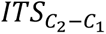 represent the interaction scores from cell type 1 to cell type 2 and from cell type 2 to cell type 1, respectively. 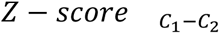 and 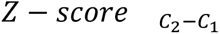 denote the standardized interaction scores from cell type 1 to cell type 2 and from cell type 2 to cell type 1, respectively. *C_1_* refers to the first interacting cell type, and *C_2_* refers to the second interacting cell type. 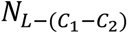 represents the number of edges between cell types *C_1_* and *C_2_*. 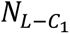 epresents the total number of edges connected to cell type *C_1_*. 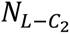 represents the total number of edges connected to cell type *C_2_*. 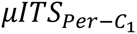 represents the mean *ITS* obtained from the permutation test, and 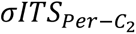 represents the variance of the *ITS* from the permutation test.

### Acquisition of Spatial Trascriptomics Data

The spatial transcriptomics data of human tonsil used in this study were obtained from publicly available resources deposited on the Zenodo platform.^47^ A preprocessed Seurat object was retrieved from the dataset, and the sample BCLL-10-T was selected for subsequent analyses.

### Clustering of Spatial Transcriptomics Data

After normalization of the Seurat object for the human tonsil sample BCLL-10-T, clustering was performed using the FindNeighbors and FindClusters functions. To correspond with the eight functional niches identified in the spatial proteomic data, the sample was divided into eight clusters. The spatial distribution of these clusters was visualized using the SpatialDimPlot function.

Differential Expression and Enrichment Analysis of Spatial Transcriptomic Subclusters Based on the seurat_clusters classification information within the Seurat object of the human tonsil spatial transcriptomics dataset, differential gene expression analysis was performed. Genes with an adjusted p-value < 0.05 and meeting the corresponding avg_log2FC thresholds for each analysis were identified as significantly differentially expressed. Subsequently, GO enrichment analysis (ontology = “BP”) was conducted to determine biological processes enriched among these genes, and the resulting pathways were categorized into distinct functional classes. Finally, functional marker genes identified from single-cell transcriptomic analysis were mapped onto the spatial transcriptomics data, and their spatial expression patterns were visualized using the SpatialFeaturePlot function.

## DATA AVAILABILITY

The original data can be obtained by contacting the corresponding author, Chengjian Zhao. All computer codes used for analyzing spatial distributions, along with further details, are available from the corresponding author upon request. The remaining data are available in the Supplemental Materials.

## ACKNOWLEDGEMENTS

This research was supported by various funding sources, including National Key Scientific and Technological Project of Ningxia Medical University (grant no. 23H1529), National Natural Science Foundation of China (grant no. 82270542), National Key Research and Development Program of China (grant no. 2023YFB330850103).

## AUTHOR CONTRIBUTIONS

Ruihan Zhou, Chaoxin Xiao and Ping Zhou contributed equally to this work. Ruihan Zhou was responsible for the conception and design of the study. Chaoxin Xiao and Ping Zhou performed the experiments. Jiaxin Liu, Xiaohong Yao, and Banglei Yin carried out the data analysis. Ouying Yan, Wanting Hou, Yulin Wang, and Huanhuan Wang contributed to the acquisition of data. Rui Zhu, Zirui Wang, and Leyi Yao provided critical revisions and helped with data interpretation. Xiaoying Li, Tongtong Xu, Fujun Cao and Na Xiao contributed to the drafting of the manuscript. Lili Jiang, Dan Cao and Ke Cheng provided technical support and supervision. Ruihan Zhou was also responsible for figure preparation. Lili Jiang supervised the project and provided critical guidance throughout the study. Chengjian Zhao oversaw the entire project and was responsible for the final approval of the manuscript.

## REFERENCES

1. Elinav, E. et al. Inflammation-induced cancer: crosstalk between tumours, immune cells and microorganisms. Nat Rev Cancer 13, 759–771, (2013).

2. Hinshaw, D. C. & Shevde, L. A. The Tumor Microenvironment Innately Modulates Cancer Progression. Cancer Res 79, 4557–4566, (2019).

3. Aliazis, K. et al. The tumor microenvironment’s role in the response to immune checkpoint blockade. Nat Cancer 6, 924–937, (2025).

4. Gajewski, T. F., Schreiber, H. & Fu, Y. X. Innate and adaptive immune cells in the tumor microenvironment. Nat Immunol 14, 1014–1022, (2013).

5. Liu, Q. et al. Cancer stem cells and their niche in cancer progression and therapy. Cancer Cell Int 23, 305, (2023).

6. Patras, L., Shaashua, L., Matei, I. & Lyden, D. Immune determinants of the pre-metastatic niche. Cancer Cell 41, 546–572, (2023).

7. Sang-Aram, C., Browaeys, R., Seurinck, R. & Saeys, Y. Unraveling cell-cell communication with NicheNet by inferring active ligands from transcriptomics data. Nat Protoc 20, 1439–1467, (2025).

8. Liu, Y. et al. Conserved spatial subtypes and cellular neighborhoods of cancer-associated fibroblasts revealed by single-cell spatial multi-omics. Cancer Cell 43, 905–924.e906, (2025).

9. Sussman, J. H. et al. Multiplexed Imaging Mass Cytometry Analysis Characterizes the Vascular Niche in Pancreatic Cancer. Cancer Res 84, 2364–2376, (2024).

10. Maestri, E. et al. Spatial proximity of tumor-immune interactions predicts patient outcome in hepatocellular carcinoma. Hepatology 79, 768–779, (2024).

11. Pi, Y. N., Guo, J. N., Lou, G. & Cui, B. B. Comprehensive analysis of prognostic immune-related genes and drug sensitivity in cervical cancer. Cancer Cell Int 21, 639, (2021).

12. Karnell, J. L., Rieder, S. A., Ettinger, R. & Kolbeck, R. Targeting the CD40-CD40L pathway in autoimmune diseases: Humoral immunity and beyond. Adv Drug Deliv Rev 141, 92–103, (2019).

13. Jiang, X. et al. Role of the tumor microenvironment in PD-L1/PD-1-mediated tumor immune escape. Mol Cancer 18, 10, (2019).

14. Kaufman, J., Sime, P. J. & Phipps, R. P. Expression of CD154 (CD40 ligand) by human lung fibroblasts: differential regulation by IFN-gamma and IL-13, and implications for fibrosis. J Immunol 172, 1862–1871, (2004).

15. Garris, C. S. et al. Successful Anti-PD-1 Cancer Immunotherapy Requires T Cell-Dendritic Cell Crosstalk Involving the Cytokines IFN-γ and IL-12. Immunity 49, 1148–1161.e1147, (2018).

16. Han, J., Wu, M. & Liu, Z. Dysregulation in IFN-γ signaling and response: the barricade to tumor immunotherapy. Front Immunol 14, 1190333, (2023).

17. Chen, H. et al. Integrative spatial analysis reveals tumor heterogeneity and immune colony niche related to clinical outcomes in small cell lung cancer. Cancer Cell 43, 519–536.e515, (2025).

18. Liu, C. C. et al. Multiplexed Ion Beam Imaging: Insights into Pathobiology. Annu Rev Pathol 17, 403–423, (2022).

19. Cheng, K. et al. Spatial interactions of immune cells as potential predictors to efficacy of toripalimab plus chemotherapy in locally advanced or metastatic pancreatic ductal adenocarcinoma: a phase Ib/II trial. Signal Transduct Target Ther 9, 321, (2024).

20. Peng, H. et al. Multiplex immunofluorescence and single-cell transcriptomic profiling reveal the spatial cell interaction networks in the non-small cell lung cancer microenvironment. Clin Transl Med 13, e1155, (2023).

21. Kleinberg, G., Wang, S., Comellas, E., Monaghan, J. R. & Shefelbine, S. J. Usability of deep learning pipelines for 3D nuclei identification with Stardist and Cellpose. Cells Dev 172, 203806, (2022).

22. Burgermeister, S. et al. Unsupervised Clustering of Cell Populations in Germinal Centers Using Multiplexed Immunofluorescence. Biology (Basel) 14, (2025).

23. Buckup, M., et al. Multiparametric cellular and spatial organization in cancer tissue lesions with a streamlined pipeline. Nat Biomed Eng, (2025).

24. Wang, H. et al. CCF-GNN: A Unified Model Aggregating Appearance, Microenvironment, and Topology for Pathology Image Classification. IEEE Trans Med Imaging 42, 3179–3193, (2023).

25. Liu, Y. et al. Identification of a tumour immune barrier in the HCC microenvironment that determines the efficacy of immunotherapy. J Hepatol 78, 770–782, (2023).

26. Gouin, K. H., 3rd et al. An N-Cadherin 2 expressing epithelial cell subpopulation predicts response to surgery, chemotherapy and immunotherapy in bladder cancer. Nat Commun 12, 4906, (2021).

27. Karimi, E. et al. Single-cell spatial immune landscapes of primary and metastatic brain tumours. Nature 614, 555–563, (2023).

28. Galeano Niño, J. L., et al. Effect of the intratumoral microbiota on spatial and cellular heterogeneity in cancer. Nature 611, 810–817, (2022).

29. Jing, S. Y. et al. Spatial multiomics reveals a subpopulation of fibroblasts associated with cancer stemness in human hepatocellular carcinoma. Genome Med 16, 98, (2024).

30. Choi, J. et al. Spatial organization of the mouse retina at single cell resolution by MERFISH. Nat Commun 14, 4929, (2023).

31. Fang, R. et al. Conservation and divergence of cortical cell organization in human and mouse revealed by MERFISH. Science 377, 56–62, (2022).

32. Liu, Q. et al. Single-cell, single-nucleus and xenium-based spatial transcriptomics analyses reveal inflammatory activation and altered cell interactions in the hippocampus in mice with temporal lobe epilepsy. Biomark Res 12, 103, (2024).

33. Zhu, J. et al. Mapping cellular interactions from spatially resolved transcriptomics data. Nat Methods 21, 1830–1842, (2024).

34. Cilento, M. A., Sweeney, C. J. & Butler, L. M. Spatial transcriptomics in cancer research and potential clinical impact: a narrative review. J Cancer Res Clin Oncol 150, 296, (2024).

35. Nave, H., Gebert, A. & Pabst, R. Morphology and immunology of the human palatine tonsil. Anat Embryol (Berl) 204, 367–373, (2001).

36. Liu, Y. et al. Effects of tonsillectomy and adenoidectomy on the immune system. Heliyon 10, e32116, (2024).

37. Maynard, K. R. et al. Transcriptome-scale spatial gene expression in the human dorsolateral prefrontal cortex. Nat Neurosci 24, 425–436, (2021).

38. Zhang, M. et al. Spatially resolved cell atlas of the mouse primary motor cortex by MERFISH. Nature 598, 137–143, (2021).

39. He, S. et al. High-plex imaging of RNA and proteins at subcellular resolution in fixed tissue by spatial molecular imaging. Nat Biotechnol 40, 1794–1806, (2022).

40. Walsh, L. A. & Quail, D. F. Decoding the tumor microenvironment with spatial technologies. Nat Immunol 24, 1982–1993, (2023).

41. Safieddine, A. et al. HT-smFISH: a cost-effective and flexible workflow for high-throughput single-molecule RNA imaging. Nat Protoc 18, 157–187, (2023).

42. Coleman, K., Hu, J., Schroeder, A., Lee, E. B. & Li, M. SpaDecon: cell-type deconvolution in spatial transcriptomics with semi-supervised learning. Commun Biol 6, 378, (2023).

43. Cable, D. M. et al. Robust decomposition of cell type mixtures in spatial transcriptomics. Nat Biotechnol 40, 517–526, (2022).

44. Xiao, C. et al. Pipeline for Assessing Tumor Immune Status Using Superplex Immunostaining and Spatial Immune Interaction Analysis. 2024.2008.2023.609368, (2024).

45. Bankhead, P. et al. QuPath: Open source software for digital pathology image analysis. Sci Rep 7, 16878, (2017).

46. Stevens, M. et al. StarDist Image Segmentation Improves Circulating Tumor Cell Detection. Cancers (Basel) 14, (2022).

47. Massoni-Badosa, R. et al. An atlas of cells in the human tonsil. Immunity 57, 379–399.e318, (2024).

48. Hu, J. et al. Tumor microenvironment remodeling after neoadjuvant immunotherapy in non-small cell lung cancer revealed by single-cell RNA sequencing. Genome Med 15, 14, (2023).

49. Wang, H. et al. SOAPy: a Python package to dissect spatial architecture, dynamics, and communication. Genome Biol 26, 80, (2025).

50. Lemaitre, L. et al. Spatial analysis reveals targetable macrophage-mediated mechanisms of immune evasion in hepatocellular carcinoma minimal residual disease. Nat Cancer 5, 1534–1556, (2024).

