## Supplementary Materials for "Multi-Omics Reveals Activated Fibroblasts with Dual Functional Roles in Repair and Negative Barrier in the CD8^+^ T Cell Cytotoxic Niche for Lung Cancer Neoadjuvant Therapy"

**This PDF file includes:**

Fig. S1 to S7

Tables S1 to S2

Fig. S1

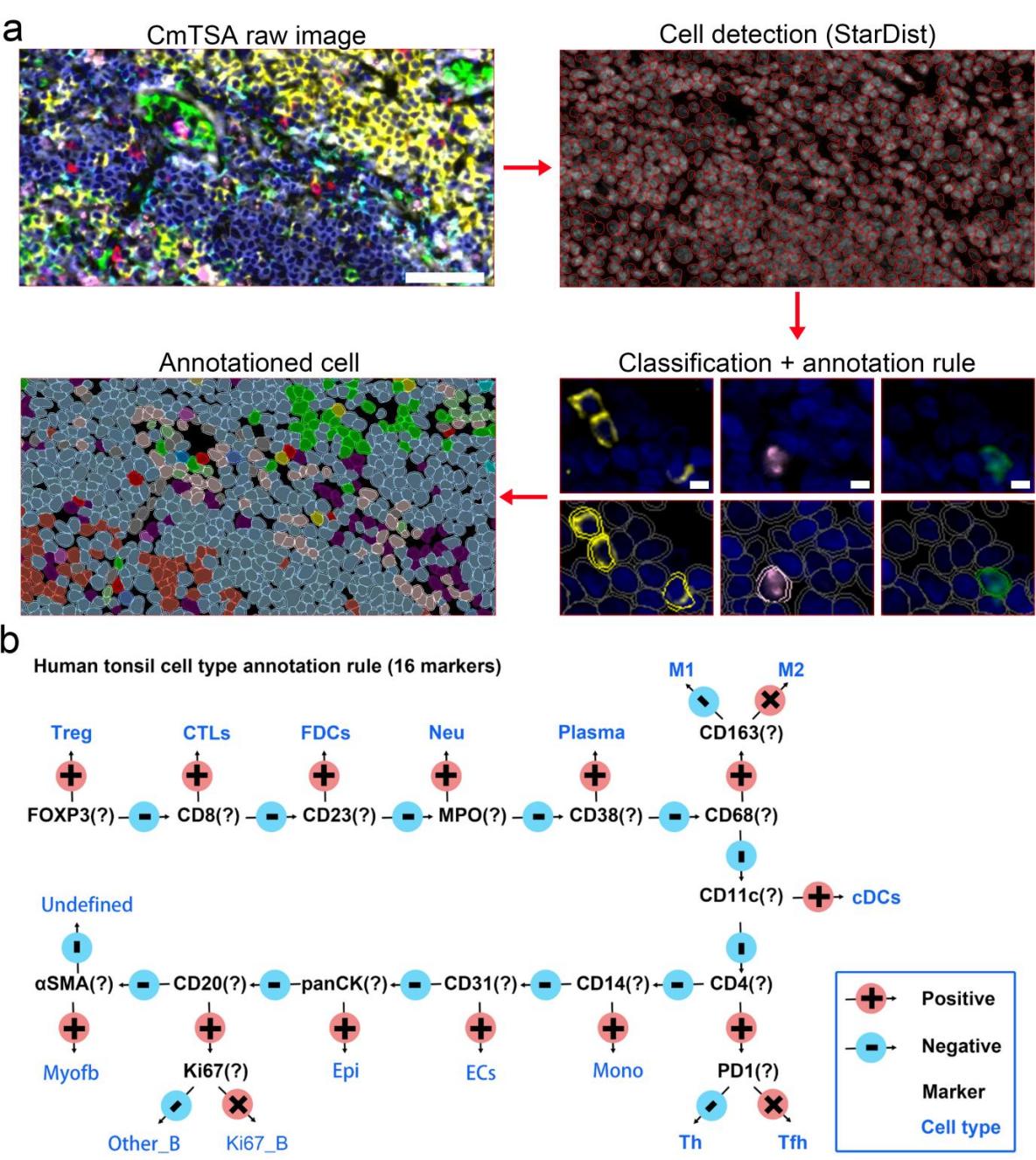

**Fig. S1 Cell type annotation strategy for the human tonsil CmTSA stained image shown in Fig. 1c**

**a** Workflow for cell annotation. CmTSA-stained images were processed in QuPath using a pretrained StarDist model for nuclear segmentation. Positive cell classes were identified by applying predefined intensity thresholds, and cell labels were assigned automatically using an Annotation Rule. The scale bar in the upper-left image represents 50 μm, and the scale bar in the lower-right image represents 5 μm.

**b** Annotation rules were defined based on canonical lineage markers, expected population abundances, and staining quality control.

Fig. S2

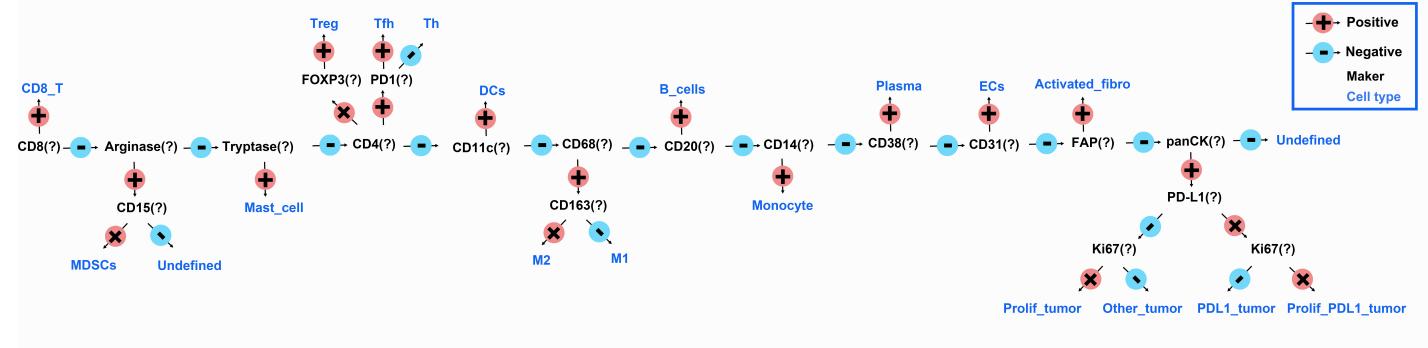

**Fig. S2 Cell type annotation strategy for the Lung cancer CmTSA stained images**

Annotation rules were defined based on canonical lineage markers, expected population abundances, and staining quality control.

**Fig. S3**

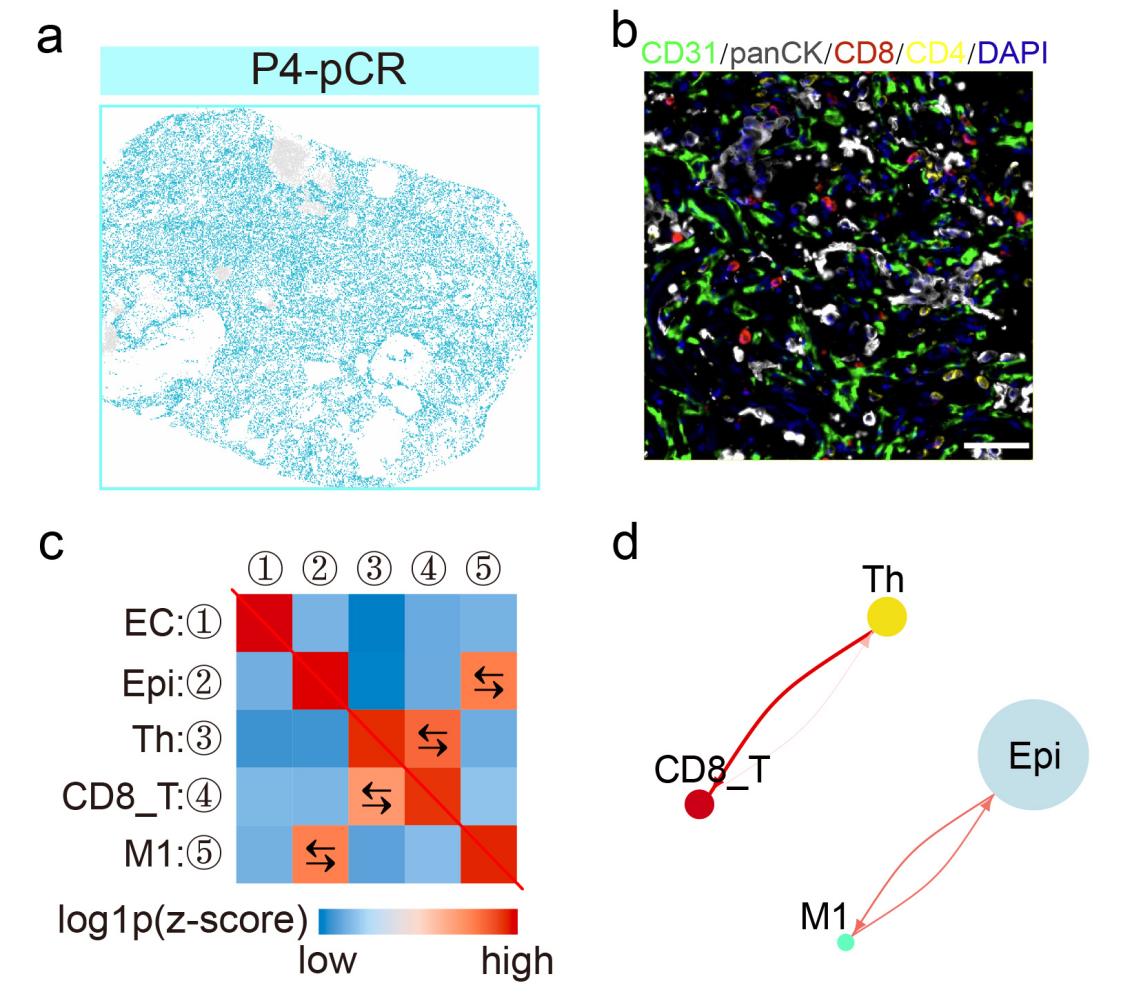

Fig. S3 Spatial map and structural organization of the P4-pCR ascularization and tissue repair FN.

**a** Spatial map of the Vascularization and Tissue Repair functional niche (FN6, blue) in this sample.

**b** Representative in situ cell images of the FN6 region in this sample, the scale bar is 50 μm.

**c** Heatmap of directional spatial interactions among different cell types within FN6, shown as log-transformed Z-scores. Color intensity reflects interaction strength, with red indicating positive interactions and blue indicating no significant interaction or spatial avoidance. Arrows denote the directionality of interactions.

**d** Cell-cell interaction network diagram, where node size represents the proportion of each cell type within FN6.

**Fig. S4**

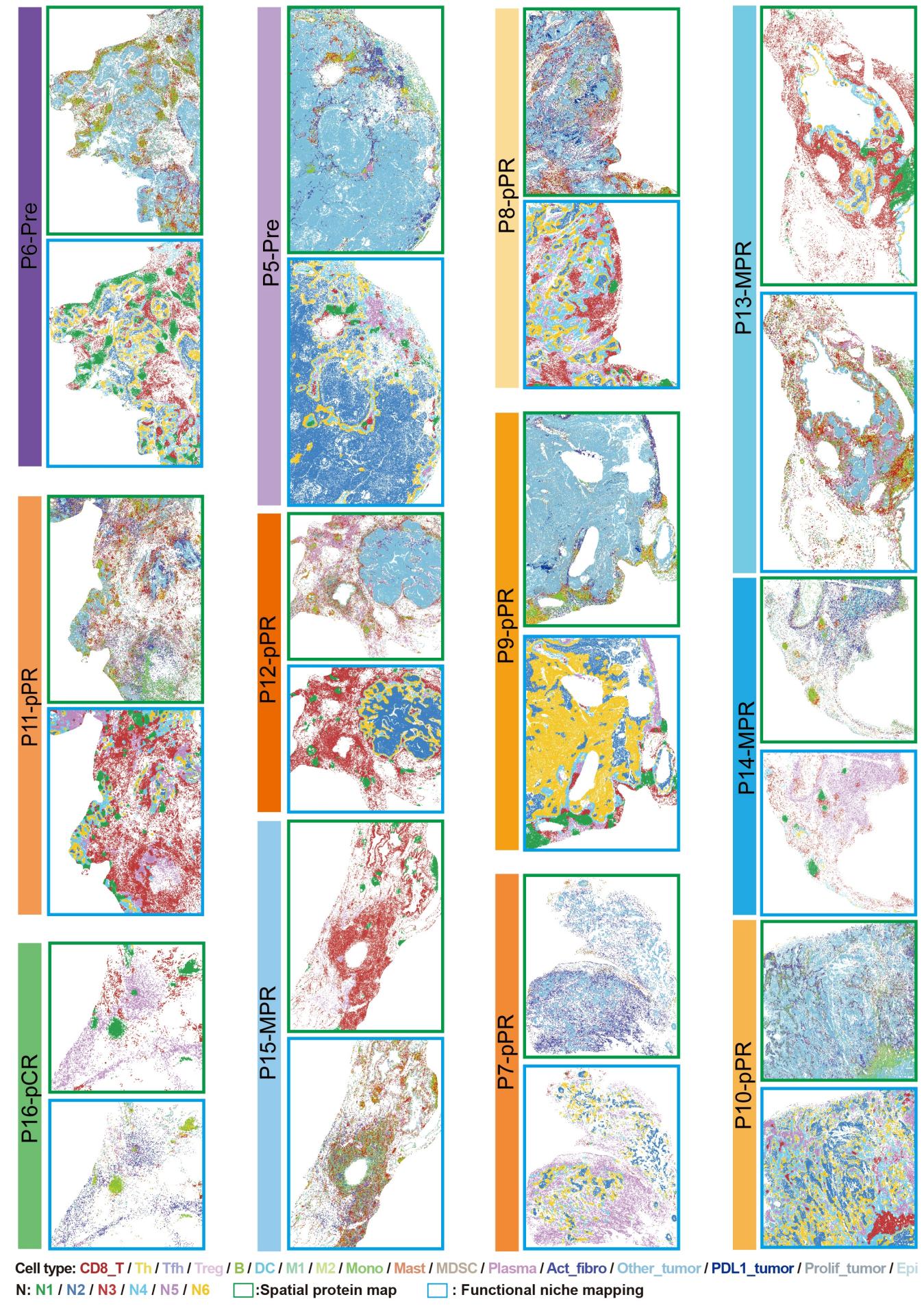

Fig. S4 Spatial proteomic maps and functional niche maps of the 12 samples in the validation group

Color scheme: Purple tones represent the pre-treatment group; orange tones represent the pPR group; blue tones represent the MPR group; and green tones represent the pCR group. Green boxes denote the spatial proteomic maps, and blue boxes denote the functional niche maps.

Fig. S5

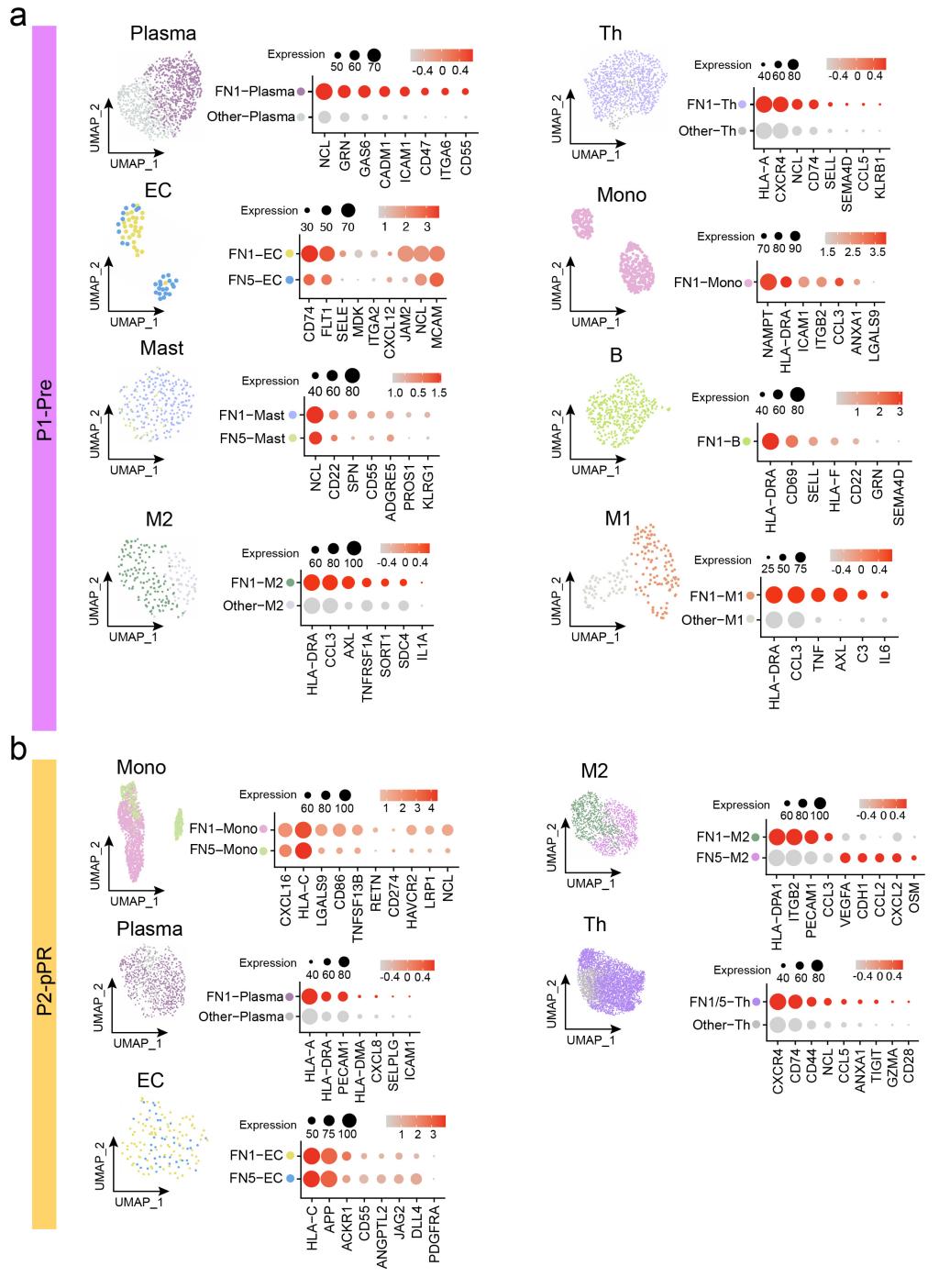

**Fig. S5 Gene expression features of CD8_T cytotoxic FNs in P1-Pre and P2-pPR samples**

**a** P1-Pre: UMAP plots show the spatial distribution of each cell type, and bubble plots display the expression levels of FN-related genes.

**b** As in panel (**a**), with the corresponding results shown for the P2-pPR sample.

Fig. S6

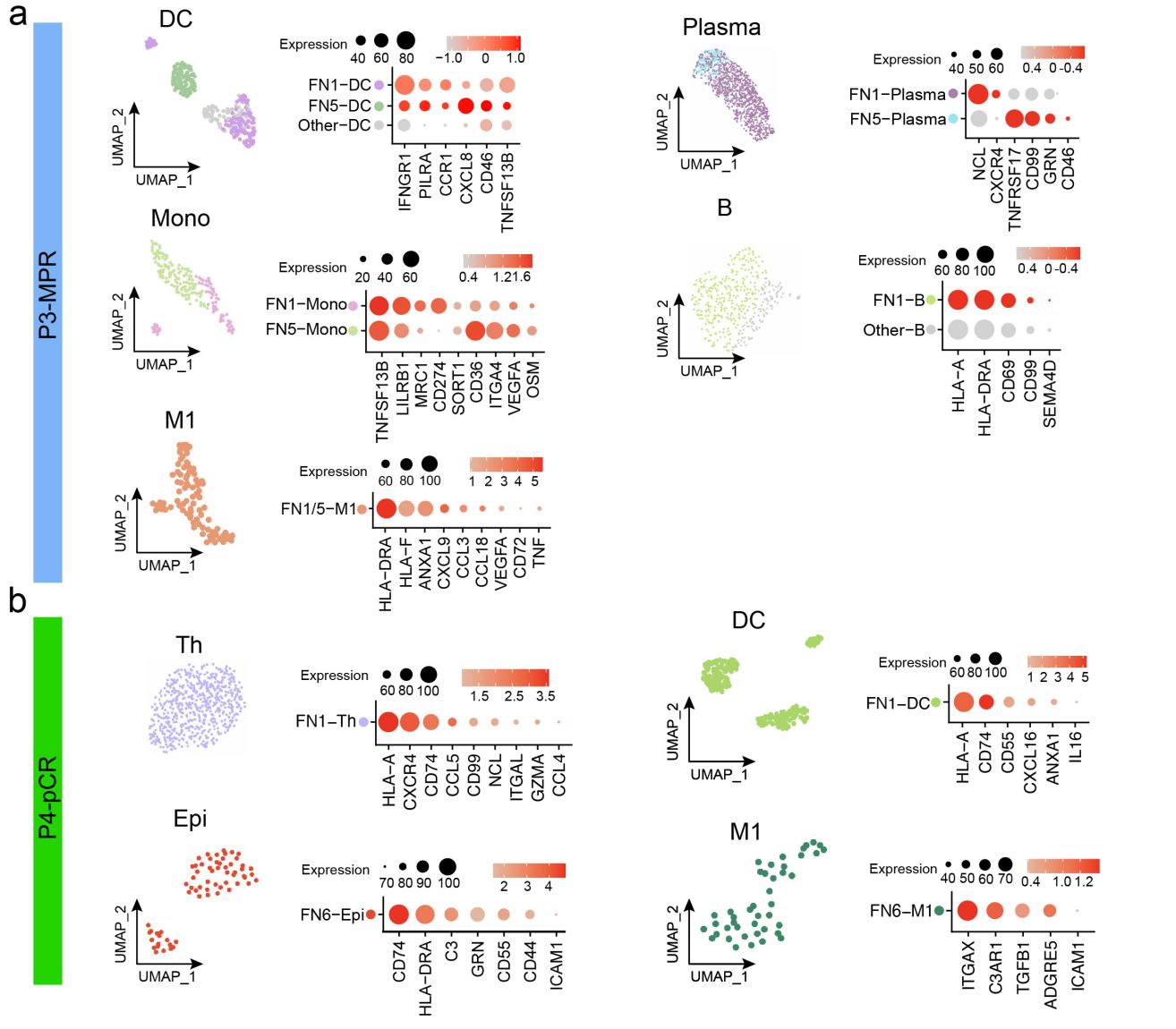

**Fig. S6 Gene expression features of CD8_T cytotoxic FNs in P3-MPR and P4-pCR samples**

**a** P3-MPR: UMAP plots show the spatial distribution of each cell type, and bubble plots display the expression levels of FN-related genes.

**b** As in panel (**a**), with the corresponding results shown for the P4-pCR sample.

**Fig. S7**

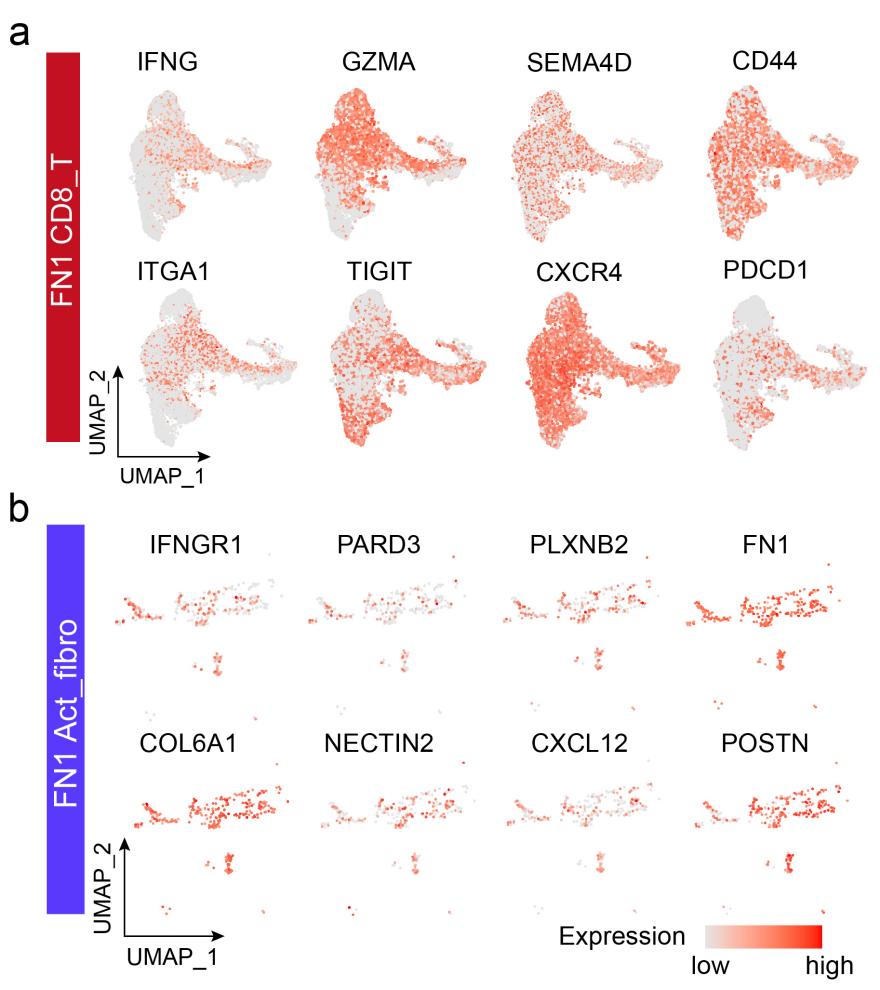

**Fig. S7 Feature plots of cytotoxic CD8_T and Act_fibro within the post-treatment CD8_T cytotoxic FNs**

**a** Gene expression intensity is visualized on spatial UMAPs, highlighting functional differences and interaction-associated signaling features of cytotoxic CD8_T cells in the niche.

**b** As in panel (**a**), for Act_fibro.

Table S1. Clinical information of NSCLC samples used in the study

| **Patient_ID** | **Sex** | **Age** | **T stage** | **N stage** | **M stage** | **Clinical tage** | **Chemotherapy** | **PD1_name** | **Response** |
| --- | --- | --- | --- | --- | --- | --- | --- | --- | --- |
| P1-Pre | Male | 65 | T1 | N2 | M0 | IIB | / | / | BL |
| P2-pPR | Male | 59 | T2a | N2 | M0 | IIIa | NP + CBP | Pembrolizumab | pPR |
| P3-pCR | Male | 65 | T4 | N2 | M0 | IIIB | NP + CBP | Terlilizumab | pCR |
| P3-MPR | Male | 59 | T3 | N2 | M0 | IIIB | NP + CBP | Terlilizumab | MPR |
| P5-Pre | Male | 65 | T2 | N2 | M0 | IIIA | / | / | BL |
| P6-Pre | Male | 67 | T1c | N2 | M0 | IIIA | / | / | BL |
| P7-pPR | Male | 55 | T3 | N1 | M0 | IIIA | DTX + DDP | Pembrolizumab | pPR |
| P8-pPR | Male | 57 | T4 | N1 | M0 | IIIA | NP + CBP | Pembrolizumab | pPR |
| P9-pPR | Male | 65 | T3 | N2 | M0 | IIIA | PTX + CBP | Toripalimab | pPR |
| P10-pPR | Male | 65 | T4 | N0 | M0 | IIIA | PTX + NDP | Sintilimab | pPR |
| P11-pPR | Male | 67 | T2a | N3 | M0 | IIIB | NP + CBP | Terlilizumab | pPR |
| P12-pPR | Male | 70 | T3 | N2 | M0 | IIIB | NP + CBP | Camrelizumab | pPR |
| P13-MPR | Male | 63 | T3 | N2 | M1b | IVa | NP + DDP | Camrelizumab | MPR |
| P14-MPR | Male | 59 | T4 | N0 | M0 | IIIA | NP + CBP | Terlilizumab | MPR |
| P15-MPR | Male | 57 | T2 | N2 | M0 | IIIA | PEM + CBP | Camrelizumab | MPR |
| P16-pCR | Male | 60 | T4 | N2 | M0 | IIIB | NP + CBP | Nivolumab | pCR |

**Respons**e: Response Evaluation Criteria In Solid Tumours

pCR: pathologic complete response; MPR: major pathologic response; pPR: partial pathological response; BL:Baseline

**Chemotherapy**

PTX:Paclitaxel; NP:nab-Paclitaxel; NDP:Nedaplatin; CBP:Carboplatin; DDP:Cisplatin

PEM:Pemetrexed; DTX:Docetaxel

**Table S2. Antibody information**

| Antibody | Brand | Catalog# | Source **s**pecies | Dilution rate |
| --- | --- | --- | --- | --- |
| aSMA | HUABIO | IRS094RT | Rat | 1:100 |
| CD11c | HUABIO | IRS103RT | Rat | 1:100 |
| CD14 | HUABIO | IRS106RT | Rat | 1:100 |
| PD-L1 | HUABIO | IRS193RT | Rat | 1: 100 |
| Tryptase | HUABIO | IRS209RT | Rat | 1:100 |
| CD38 | HUABIO | IRS119RT | Rat | 1:100 |
| CD68 | HUABIO | IRS004 | Mouse | 1:100 |
| CD8 | HUABIO | IRS007 | Mouse | 1: 100 |
| MPO | HUABIO | IRS173MS | Mouse | 1:100 |
| panCK | HUABIO | IRS010 | Mouse | 1:100 |
| PD-1 | HUABIO | IRS190MS | Mouse | 1: 100 |
| CD20 | HUABIO | IRS112MS | Mouse | 1:100 |
| CD31 | HUABIO | IRS023 | Mouse | 1:100 |
| CD4 | HUABIO | IRS004 | Rabbit | 1:100 |
| FAP | HUABIO | IRS289RB | Rabbit | 1: 100 |
| FOXP3 | HUABIO | IRS008 | Rabbit | 1: 100 |
| Ki67 | HUABIO | IRS057 | Rabbit | 1:100 |
| CD23 | HUABIO | IRS021 | Rabbit | 1:100 |
| CD163 | HUABIO | IRS002 | Rabbit | 1:100 |
| Arginase1 | HUABIO | IRS081RB | Rabbit | 1:100 |
| CD15 | HUABIO | IRS107RB | Rabbit | 1:100 |
